# How suitable are clustering methods for functional annotation of proteins?

**DOI:** 10.1101/2024.12.26.630370

**Authors:** Rakesh Busi, Pranav Machingal, Nandyala Hemachandra, Petety V. Balaji

## Abstract

The advent of affordable high-throughput genome sequencing has drastically expanded protein sequence databases, necessitating the development of computational tools to predict protein function from sequence data. Current methods, such as BLASTp and profile HMMs, while effective, are limited by difficulties in detecting remote homologs and uncertainties in multiple sequence alignments. To address this, we explore the use of clustering algorithms for unsupervised protein function annotation, using pseudo-amino acid composition (PAAC) as features.

In this study, we evaluated nine clustering algorithms for their ability to segregate protein sequences based on functional differences using the PAAC feature. Using intrinsic metrics, particularly the silhouette coefficient (SC), we determined the optimal number of clusters (*k*) for each algorithm. We observed that agglomerative clustering produced results resembling phylogenetic relationships; even k-means clustering, Gaussian mixture model (GMM), and spectral clustering do so, but occasionally merge datapoints from distinct original clusters at higher *k* values.

Our findings reveal that k-means clustering, GMM, and agglomerative clustering effectively segregate distinct protein functional families, but effectiveness decreases when distinguishing fine-grained functional differences. Notably, spectral clustering underperformed relative to other methods. Affinity propagation clustering, while effective in some cases, generated more clusters than expected and is prone to false positives. Overall, we find that some of the clustering algorithms are suitable for functional annotation of protein sequences using PAAC as a feature set, even when the number of ground-truth sequences is limited.

The implementation of the clustering method for protein sequences is available in the GitHub repository (https://github.com/RakeshBusi/Clustering). It provides comprehensive steps for preprocessing, feature extraction, clustering, and evaluation. All steps are presented in a Jupyter Notebook in the repository.

**Author Summary:** We are in the age of big data. It is an outcome of the development of high-throughput techniques. The resources spent to develop and deploy such techniques are considerably large. However, data by itself is not an end but a means to answer questions of relevance. Hence, the development and/or customisation of techniques that help us to interpret and utilise data are also important. In this study, we focus on customising a popular technique, namely clustering, to extract biological information from the ever-growing protein sequence database. We test the suitability of nine clustering algorithms to determine a protein’s molecular function solely based on its amino acid sequence. Based on our findings, we recommend using a combination of the four algorithms, namely, k-means, Gaussian mixture model, agglomerative, and affinity propagation. However, we note that proteins with subtle functional differences cluster together, and fine-tuning algorithms to separate such proteins requires additional experimental data.

## Introduction

Affordable high-throughput genome sequencing has led to an enormous increase in the size of protein sequence databases [1]. Knowledge of the functions of these proteins is necessary to fully realize the benefits of genome sequencing. Experimental characterization of the functions of all these proteins is impractical. This hurdle can be circumvented to a large extent by computational methods that can predict possible function(s) based primarily on amino acid sequence.

Currently, BLAST [2] (including its variants), profile HMMs [3], and position weight matrix-based motif search [4] are the computational methods of choice for function annotation. As sequences diverge, it becomes difficult to decide if two homologs are paralogs or orthologs. Clustering algorithms, being unsupervised methods, do not need labels for datapoints. This allows one to combine experimentally characterized (i.e., functionally annotated) and unannotated sequences for assigning functions to the latter. Working towards such a broad goal, our focus in this study is to assess the ability of clustering methods to segregate input protein sequences along the lines of their functional differences. Herein, we report the clustering of functionally distinct protein sequences using pseudo amino acid composition (PAAC) [5] as features.

In the current study to evaluate the effectiveness of clustering methods in protein annotation, we selected datasets containing protein sequences labelled with fine-grained functional differences. Specifically, we focus on Glycoside Hydrolases (GH), which are crucial enzymes in carbohydrate metabolism. The GH dataset was chosen for several reasons: first, it consists of manually curated protein sequences from the widely recognized CAZy database [6]; second, the GH enzymes are categorized by EC numbers (EC 3.2.1.x) [7], where the first three digits represent glycosidases that hydrolyze O- and S-glycosyl compounds, and the final digit specifies substrate specificity, indicating fine-grained functional differences; third, GH enzymes are grouped into GH families in the CAZy database, which not only reflect structural features but also reveal evolutionary relationships [8]. In addition to GH datasets, we included datasets with broader functional differences to assess the robustness of clustering methods across a wider spectrum of protein functions.

## Methods

### Datasets

We have considered a diverse range of 15 datasets, each of which contains ≥2 protein families (Table 1). Depending on the presence (or lack of) sequence, 3D structural, and/or functional similarity between protein families constituting a dataset, the 15 datasets have been divided into 5 groups. Further details of these datasets and their creation are given in the section Details related to datasets, in Supporting information S1 File.

**Table 1.** Grouping of 15 datasets into 5 groups based on shared characteristics.

| Group | Characteristics | Datasets |
| --- | --- | --- |
| Group 1 | Sequence - Dissimilar | Lysozyme CGCh (3 families; 123 sequences) |
|  | 3D structure - Different fold | Xylanase (2 families; 699 sequences) |
|  | Function - Similar | Chitinase (2 families; 656 sequences) |
| Group 2 | Sequence - Similar | Lysozyme CaLA (2 families; 74 sequences) |
|  | 3D structure - Same fold | Protease (2 families; 83 sequences) |
|  | Function - Dissimilar | Ferredoxin (2 families; 16 sequences) |
| Group 3 | Sequence - Dissimilar | β-Glucosidase (2 families; 552 sequences) |
|  | 3D structure - Same fold | Lysozyme (3 families; 213 sequences) |
| | Function - Similar | β-Galactosidase (3 families; 495 sequences) |
| Group 4 | Sequence - Dissimilar | GH2 (2 families; 202 sequences) |
|  | 3D structure - Same fold | GH3 (2 families; 303 sequences) |
|  | Function - Dissimilar | GH5 (2 families; 469 sequences) |
| Group 5 | Sequence - Dissimilar | SNARE (2 families; 118 sequences) |
|  | 3D structure - Different fold | GPCR1 (2 families; 3604 sequences) |
|  | Function - Dissimilar | GPCR2 (2 families; 3558 sequences) |

Characteristics considered include sequence, 3D structure, and function. Each group contains various combinations of similar and dissimilar characteristics. The table lists the datasets in each group, along with the number of families in each dataset and the total number of sequences.

Group 1 consists of Lysozyme CGCh, Xylanase, and Chitinase datasets. Protein families in these datasets perform the same function, i.e., sequences have the same EC number.

Group 2 consists of Lysozyme CaLA, Protease, and Ferredoxin datasets. Protein families in these datasets share sequence and 3D structure similarity but differ in their function. The difference is fine-grained in the case of the Protease dataset because chymotrypsin and trypsin catalyze the same reaction (hydrolases of the peptide bond) but differ in their substrate specificity. Lysozyme C and *α*-Lactalbumin of the Lysozyme CaLA dataset, and FDX1 and FDX2 of the Ferredoxin dataset perform mutually exclusive functions [9].

Group 3 consists of *β*-Glucosidase, Lysozyme, and *β*-Galactosidase datasets. Protein families in these datasets perform the same function, i.e., sequences have the same EC number.

Group 4 consists of GH2, GH3, and GH5 datasets, which are distinct from those in other groups: sequences from all three datasets have the same first 3 digits of EC number, i.e., 3.2.1.-. They differ in their substrate specificities, leading to differences in the 4th digit of the EC number.

Group 5 consists of SNARE, GPCR1, and GPCR2 datasets comprising proteins with no catalytic activity (no EC number). The (i) SNARE and non-SNARE in the SNARE dataset, (ii) GPCR and non-GPCR(TM) in the GPCR1 dataset, and (iii) GPCR and non-GPCR(no TM) in the GPCR2 dataset do not share any sequence, structure, or function similarities. Proteins belonging to non-SNARE do not constitute a protein family, i.e., not homologous to each other, but non-SNARE is referred to as a protein family along with all others for ease of communication; this is true for non-GPCR(TM) and non-GPCR(no TM) also.

## Clustering methods

We have assessed nine clustering algorithms provided in the scikit-learn platform [10] using pseudo amino acid composition (PAAC) [5] as features. PAAC was chosen because it captures both amino acid composition and sequence order correlation, which are crucial for determining a molecular function. The quality of clustering was assessed using (i) the silhouette coefficient, an intrinsic metric, and (ii) the Fowlkes-Mallows index, an extrinsic metric computed using database-assigned labels [11]. Further details are given in the sections Feature engineering, and Details related to clustering algorithms, in Supporting information S1 File.

## Results

### Determining the optimal number of clusters for a given dataset

The number of clusters (*k*) to be formed by input data is user-defined for k-means, GMM, agglomerative, and spectral clustering. The optimal number of clusters for each clustering algorithm was determined by evaluating clustering quality using the silhouette coefficient (SC) value, with *k* ranging from 2 to 10. A higher SC value indicates better separation between clusters. Therefore, for each algorithm, we selected the *k* value corresponding to the highest SC, as shown in Table 2. For each dataset, among the four optimal *k* values (one for each of the four algorithms), the clustering result corresponding to the *k* with the highest Fowlkes-Mallows index (FMI) was chosen as the final result. An analysis of these clustering results for each group is explained below and shown as contingency matrices. The contingency matrices of Lysozyme CGCh, *β*-Galactosidase, and GH3 datasets are shown in Table 3 while other contingency matrices are shown in Tables 4 and 5, in Supporting information S1 File.

**Table 2.** Optimal number of clusters and highest silhouette coefficient for four clustering algorithms.

| Group | Dataset | K-means |  | GMM |  | Agglomerative |  | Spectral |  |
| --- | --- | --- | --- | --- | --- | --- | --- | --- | --- |
| | | $k$ | $SC_{max}$ | $k$ | $SC_{max}$ | $k$ | $SC_{max}$ | $k$ | $SC_{max}$ |
| Group 1 | Lysozyme CGCh | 2 | 0.35 | 2 | 0.35 | 2 | 0.34 | 2 | 0.31 |
|  | Xylanase | 2 | 0.36 | 2 | 0.35 | 2 | 0.35 | 2 | 0.35 |
|  | Chitinase | 3 | 0.18 | 3 | 0.18 | 3 | 0.18 | 2 | 0.12 |
| Group 2 | Lysozyme CaLA | 2 | 0.37 | 2 | 0.37 | 2 | 0.37 | 2 | 0.37 |
|  | Protease | 2 | 0.35 | 2 | 0.35 | 2 | 0.38 | 2 | 0.38 |
|  | Ferredoxin | 6 | 0.30 | 6 | 0.30 | 5 | 0.32 | 2 | 0.22 |
| Group 3 | β-Glucosidase | 2 | 0.24 | 2 | 0.21 | 3 | 0.24 | 10 | 0.14 |
|  | Lysozymes | 10 | 0.24 | 10 | 0.24 | 10 | 0.27 | 10 | 0.21 |
| | β-Galactosidase | 3 | 0.33 | 3 | 0.33 | 3 | 0.33 | 8 | 0.15 |
| Group 4 | GH2 | 10 | 0.35 | 10 | 0.35 | 10 | 0.38 | 10 | 0.36 |
|  | GH3 | 3 | 0.25 | 3 | 0.25 | 3 | 0.24 | 2 | 0.26 |
|  | GH5 | 2 | 0.25 | 2 | 0.24 | 2 | 0.21 | 2 | 0.23 |
| Group 5 | SNARE | 2 | 0.14 | 2 | 0.14 | 2 | 0.09 | 2 | 0.13 |
|  | GPCR1 | 2 | 0.15 | 2 | 0.34 | 2 | 0.12 | 2 | 0.14 |
|  | GPCR2 | 2 | 0.22 | 2 | 0.28 | 2 | 0.20 | 2 | 0.21 |

The table shows the optimal number of clusters ( $k$ ) and corresponding highest silhouette coefficient ( $SC_{max}$ ) for k-means clustering, Gaussian mixture model (GMM), agglomerative, and spectral clustering across 15 datasets in 5 groups. Optimal $k$ was selected from a range of 2–10 based on the highest SC value, where higher SC indicates better cluster separation.

**Table 3.**
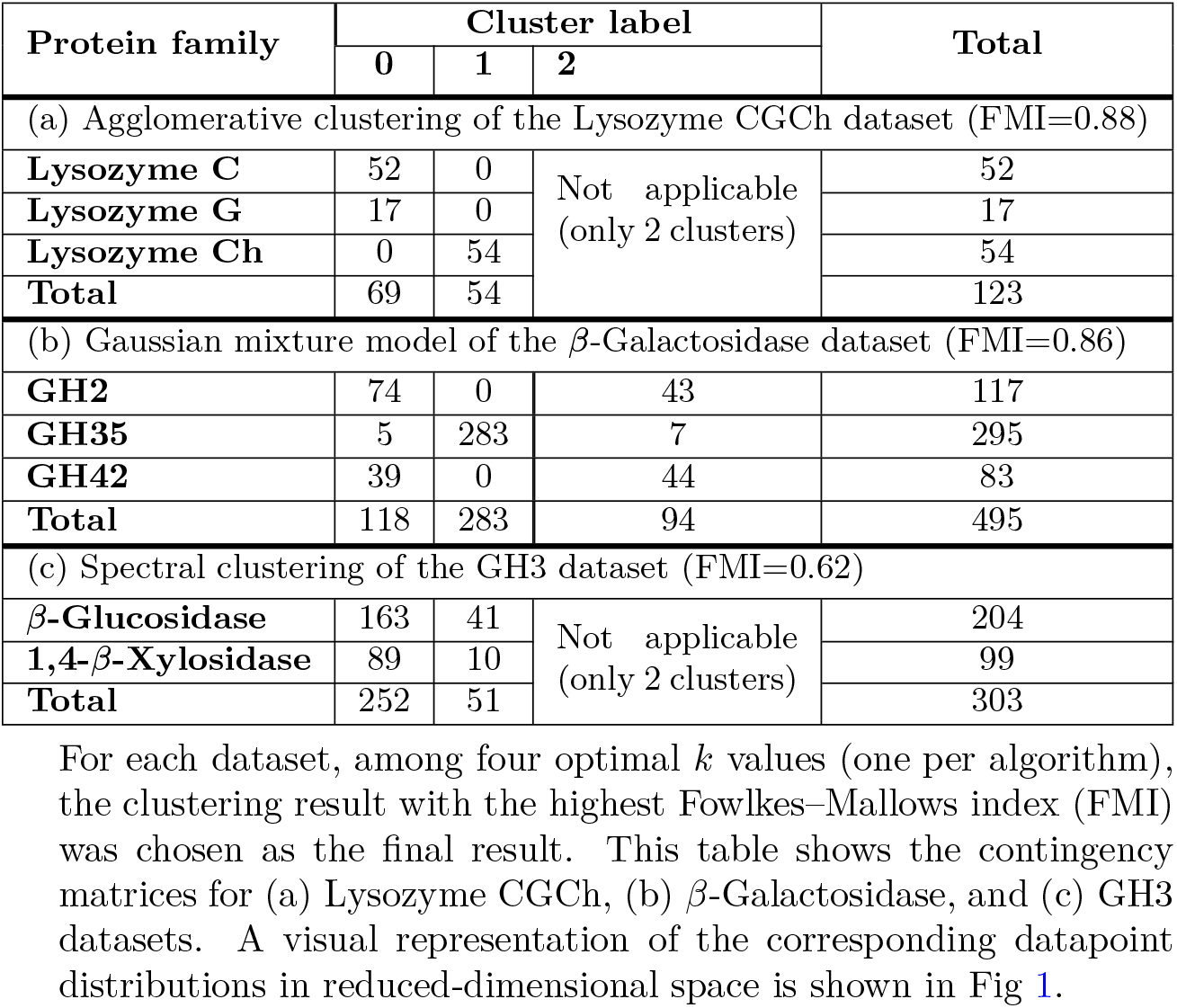
Contingency matrices of clustering results with the highest Fowlkes–Mallows index.

| Protein family | Cluster label |  |  | Total |
| --- | --- | --- | --- | --- |
|  | 0 | 1 | 2 |  |
| (a) Agglomerative clustering of the Lysozyme CGCh dataset (FMI=0.88) |  |  |  |  |
| Lysozyme C | 52 | 0 | Not applicable<br>(only 2 clusters) | 52 |
| Lysozyme G | 17 | 0 |  | 17 |
| Lysozyme Ch | 0 | 54 |  | 54 |
| Total | 69 | 54 |  | 123 |
| (b) Gaussian mixture model of the β-Galactosidase dataset (FMI=0.86) | | | | |
| GH2 | 74 | 0 | 43 | 117 |
| GH35 | 5 | 283 | 7 | 295 |
| GH42 | 39 | 0 | 44 | 83 |
| Total | 118 | 283 | 94 | 495 |
| (c) Spectral clustering of the GH3 dataset (FMI=0.62) |  |  |  |  |
| β-Glucosidase | 163 | 41 | Not applicable<br>(only 2 clusters) | 204 |
| 1,4-β-Xylosidase | 89 | 10 | | 99 |
| Total | 252 | 51 |  | 303 |

**Table 4.**
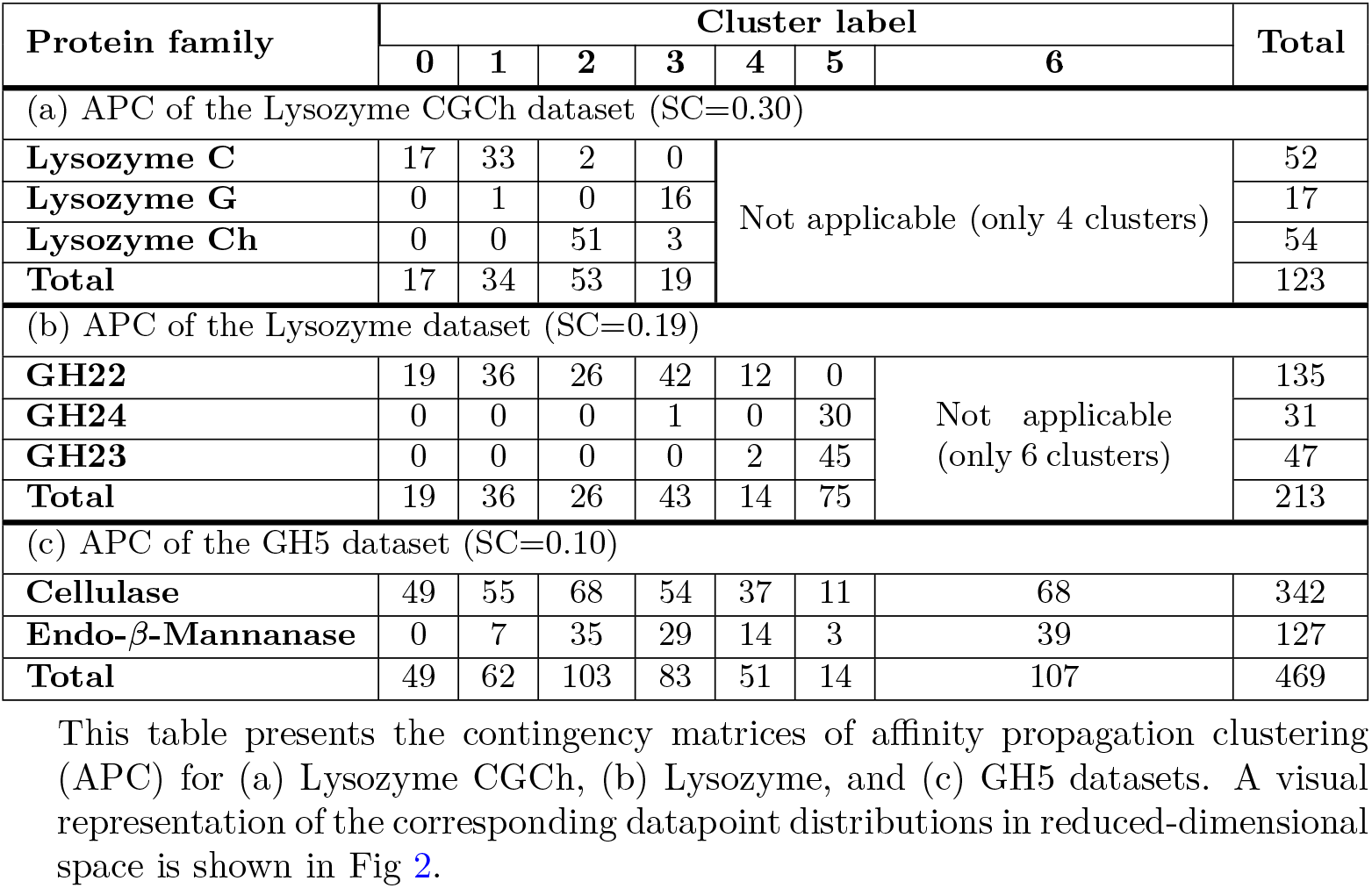
Contingency matrices of affinity propagation clustering results.

**Fig 1.**
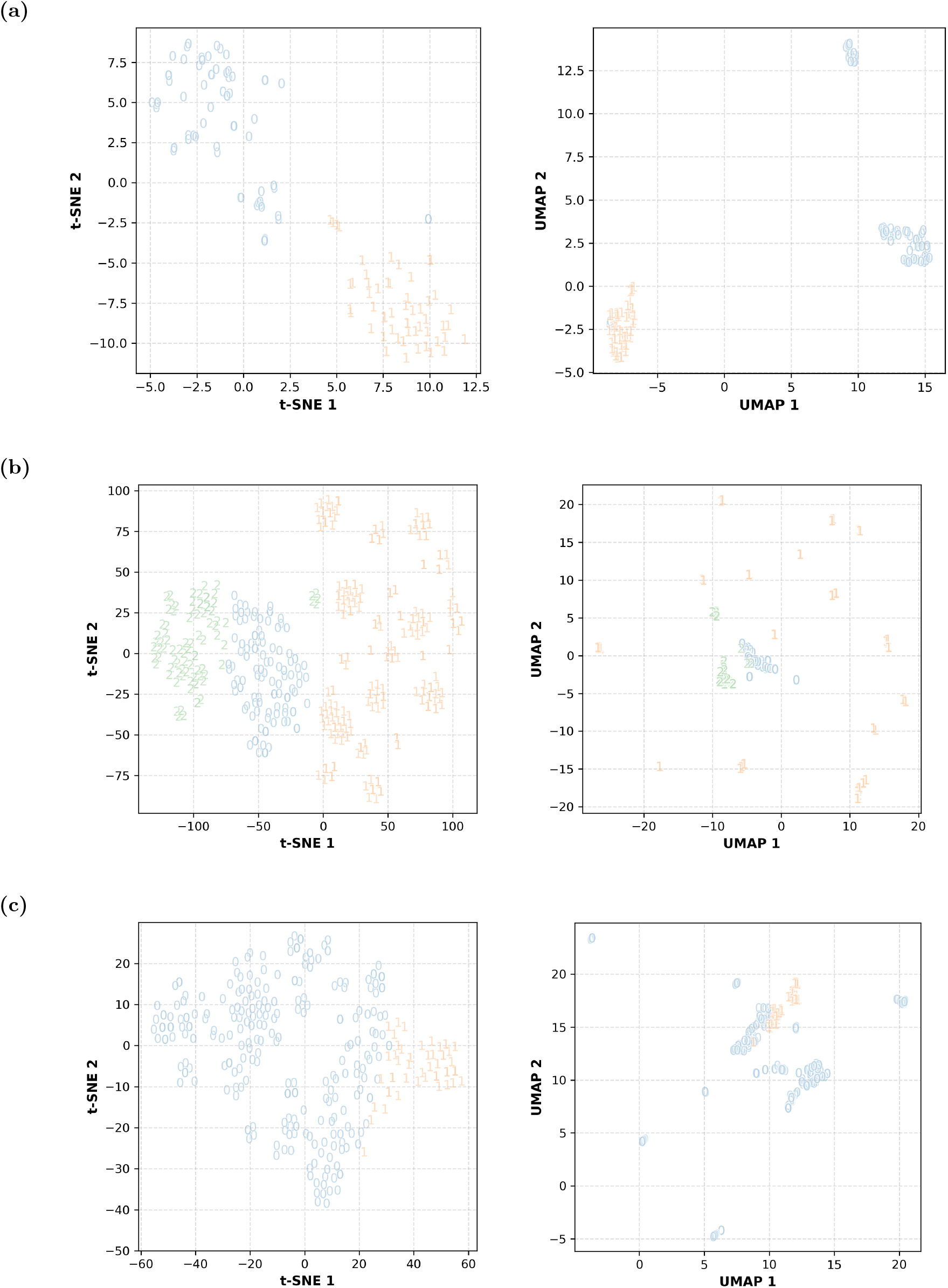
Distribution of datapoints in reduced-dimensional space for selected clustering results. The figure shows the distribution of datapoints in various clusters corresponding to the results in Table 3: (a) Lysozyme CGCh; agglomerative, (b) *β*-Galactosidase; GMM, and (c) GH3; spectral. Cluster labels are represented by symbols in the graphs. Both t-SNE and UMAP algorithms have been used for dimensionality reduction. Similar plots for other datasets are provided in Figs 7–11 of Supporting information S1 File.

**Fig 2.**
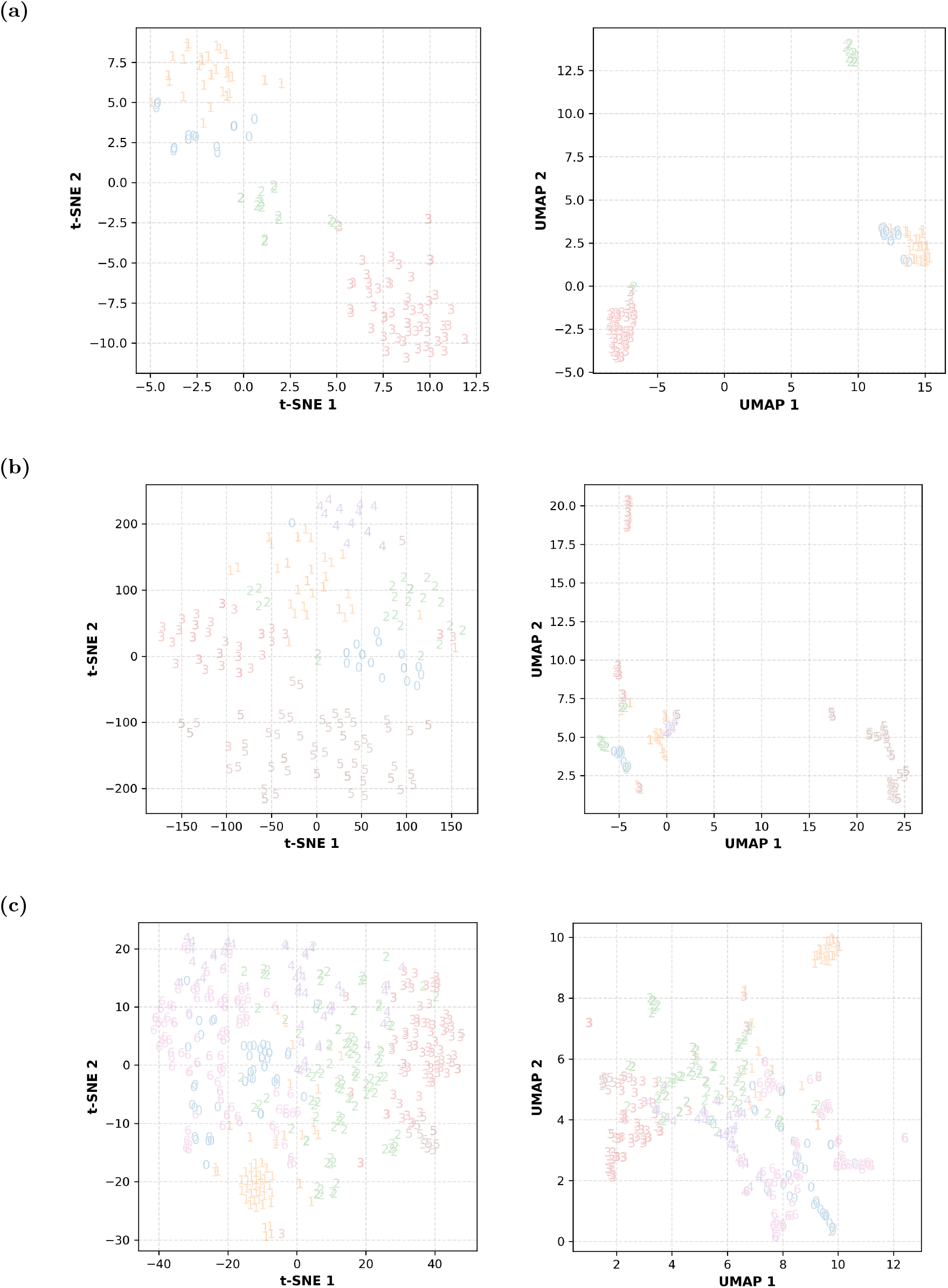
Distribution of datapoints in reduced-dimensional space for affinity propagation clustering results. The figure shows the distribution of datapoints in various clusters corresponding to the results in Table 4: APC of (a) Lysozyme CGCh, (b) Lysozyme, and (c) GH5. Cluster labels are represented by symbols in the graphs. Both t-SNE and UMAP algorithms have been used for dimensionality reduction. Similar plots for other datasets are shown in Figs 12–16 of Supporting Information S1 File.

### Group 1

Clusters formed by agglomerative/spectral clustering achieve the highest FMI for the three datasets of this group. The labels assigned by the clustering algorithm and the database agree very well in the case of the Xylanase and Chitinase datasets. In the case of the Lysozyme CGCh dataset, lysozyme C and lysozyme G sequences have been grouped into a single cluster (Table 3(a)). Hence, even though the number of families (as determined by database-assigned labels) is 3, we find only two clusters. It has been found that lysozyme C and lysozyme G share a common ancestor and hence retain the same fold. This suggests that, in this case, clustering is able to identify remote homologs.

### Group 2

All three datasets in this group have two families, and even the clustering algorithm identified two clusters. The database-assigned and cluster labels agree completely for all sequences in the Lysozyme CaLA dataset, for the majority of the sequences in the Ferredoxin dataset, but for only some sequences in the Protease dataset. In the Protease dataset, chymotrypsin and trypsin share very high similarity in their 3-D structure and function, differing only in substrate specificity. This shows that clustering algorithms do not necessarily distinguish proteins with fine-grained functional differences. In the Ferredoxin dataset, functional differences arise because of a loop region; the rest of the protein sequence and 3-D structure are the same. Clustering algorithms are able to detect this difference, and it is tempting to attribute this ability to the sequence correlation effect captured by feature engineering.

### Group 3

The agreement between the database-assigned and cluster labels is not as good as it is for the datasets in Group 1 and Group 2 (except the Protease dataset). The *β*-Glucosidase dataset consists of two families; clustering gives three clusters. Two of the three clusters have sequences predominantly belonging to either one of the two families; the third cluster has representation from both families. For the *β*-Galactosidase dataset, GH35 sequences form a distinct cluster, whereas GH2 and GH42 sequences do not separate well (Table 3(b)). The Lysozyme dataset gives 10 clusters, while it has only three families. GH22 sequences are split into multiple clusters; only four sequences from the GH23 family are part of these clusters. The separation of GH23 and GH24 sequences is not quite distinct. We have not been able to assign any possible reason for this pattern of clustering.

### Group 4

Datasets in this group are distinct from those in other groups: Sequences from all three datasets have the same first three digits of the EC number, i.e., 3.2.1.-. Sequences of the GH2 dataset form 10 clusters, even though they have two families. GH3 (Table 3(c)) and GH5 datasets also have two families, and in each of these cases, only two clusters are formed; yet there is a mismatch between cluster labels and database-assigned labels. Overall, the performance of the clustering algorithms from the viewpoint of matching with the database-assigned labels is not good.

### Group 5

Datasets in this group comprise proteins without catalytic activity (no EC number). Each of the three datasets has two categories of sequences: one category is a functional family of proteins, whereas the other consists of sequences chosen using the NOT Boolean operator to ensure that none of the sequences belong to the functional family. Because of this reason, labels are given as a *functional family* and *non-functional family*. These labels were compared with cluster labels. The two labels show good agreement with each other for the SNARE and GPCR2 datasets, but not for the GPCR1 dataset. The discrepancy in GPCR1 may be due to the presence of transmembrane domains in both GPCR and non-GPCR(TM) families, as such domains often share a similar amino acid composition, particularly being rich in apolar amino acids.

### Analysis of the number and sizes of clusters formed by affinity propagation clustering

The number of clusters into which datapoints are to be segregated is determined by the algorithm itself in the case of affinity propagation clustering (APC). The number of clusters ranges from 2 to 12, considering all 15 datasets. However, we expected 2 or 3 clusters based on database-assigned labels. SC values range from 0.15 to 0.30 except as noted here. SC values range from 0.05 to 0.11 for GH5, SNARE, GPCR1, and GPCR2, which are comparatively closer to 0, indicating overlapping clusters. The clustering results were analyzed using database-assigned labels also; a few examples are shown in Table 4, and results for all datasets can be found in Tables 6 and 7, in Supporting information S1 File. The FMI values for some of the datasets are relatively lower because the number of database-assigned labels is fewer than the number of clusters (*k*) formed, resulting in an increase in the number of false negatives.

Visual analysis of contingency matrices shows that most of the clusters formed by APC have datapoints with the same database-assigned labels in the case of Lysozyme CGCh (Table 4(a)), Xylanase, Chitinase, Lysozyme CaLA, Ferredoxin, *β*-Glucosidase, and GPCR2 datasets. This suggests that datapoints are divergent (vis-à-vis the values used for various parameters of the algorithm) in the 50-D space despite having the same database-assigned labels.

The scenario is different in the case of the remaining 8 datasets. Let us take the Lysozyme dataset as an example. There are three protein families in the Lysozyme dataset, viz., GH22, GH23, and GH24. Sequences belonging to GH22 form many clusters with hardly any datapoints from the other two families. Sequences belonging to the GH23 and GH24 families together form single clusters(Table 4(b)). This is suggestive of the proximity of GH23 and GH24 datapoints to each other in the 50-D space. From this, one can envisage three possibilities: (i) These sequences have fine-grained functional differences despite sequence similarity. (ii) PAAC is not capturing the sequence features that determine function. (iii) The database-assigned labels need revision. However, in the case of the Protease, GH3, GH5 (Table 4(c)), SNARE, and GPCR1 datasets, the database-assigned labels and cluster labels show large-scale disagreement. It appears that the database-assigned labels need revision in all these cases. Note: for the *β*-Galactosidase and GH2 datasets, APC failed to converge with the default parameters, producing degenerate cluster centers and labels that may not reflect meaningful clusters.

## Discussion

In this study, we considered nine available algorithms in the scikit-learn platform [10] to evaluate their suitability in segregating protein families reflecting functional differences. K-means, Gaussian mixture model (GMM), agglomerative, and spectral clustering were chosen because of their flexibility to allow a user-defined number of clusters, denoted as *k*. We also gave special consideration to affinity propagation clustering (APC) despite its non-user-defined *k*, as it demonstrated the ability to segregate proteins effectively.

The remaining four algorithms, mean-shift, DBSCAN, OPTICS, and BIRCH, were excluded from consideration primarily because they do not allow user-defined *k*. These algorithms either produced a single cluster or an excessive number of clusters where most data points from different classes fell into a single cluster. This outcome can be attributed to the default parameters, which may not align with the characteristics of our datasets. Furthermore, the lack of a widely accepted metric to assess intrinsic performance, especially when labels are not provided a priori, limits their applicability in this study.

To further assess clustering performance, we studied the pattern of cluster formation for values of *k* ranging from 2 to 10 for k-means, GMM, agglomerative, and spectral algorithms. Notably, agglomerative produced dendrograms resembling phylogenetic trees, which align with the phylogenetic relationships of the protein families in our dataset. Therefore, the splitting of clusters with increasing *k* values can be seen as a depiction of evolutionary divergence. However, in the case of k-means, GMM, and spectral clustering, the pattern differed; when *k* increased, some newly formed clusters contained data points from two or more original clusters. This behavior, although rare (≤10% of the data points), was occasionally observed at higher rates in 4-5 cases, with up to 30% of the data points affected.

We employed three intrinsic metrics: silhouette coefficient (SC), Calinski-Harabasz (CH) index, and Davies-Bouldin (DB) index to determine the optimal *k* for each algorithm. However, only SC proved suitable for this study, and we discarded the CH and DB indices due to their previously discussed limitations. Using SC, we identified the optimal *k* for each algorithm on each dataset, selecting the value with the highest SC score as the optimal solution for further analysis.

When comparing the optimal *k* and SC scores across the datasets, we found that the majority of cases exhibited consensus across the four clustering algorithms. Notably, in cases where consensus was absent, the discrepancy was largely due to the underperformance of spectral clustering, which often produced *k* values that differed when the SC value showed a ratio of approximately 2:3 difference from the other three algorithms for each dataset. Additionally, k-means, GMM, and agglomerative generally produced identical/similar SC values, whereas spectral clustering consistently underperformed. This raises the possibility that SC may be biased against spectral clustering or that spectral clustering is not well-suited for our datasets and features.

The suspicion of SC being algorithm-dependent prevented us from selecting a single optimal algorithm and *k* for each dataset. Instead, we carried out all four algorithms with their respective optimal *k* forward for further analysis. For the final evaluation, we utilized the highest Fowlkes-Mallows index (FMI) from these four algorithms for each dataset and plotted contingency matrices to visually assess performance.

Clustering methods (i.e., a combination of pseudo amino acid composition (PAAC) and clustering algorithms) successfully segregated the datasets in Group 1, Group 2 (except for the protease dataset), and Group 5 (except for GPCR1). Group 3 exhibited partial segregation, while Group 4 posed significant challenges. In the few exceptions where clustering methods underperformed, the discrepancies were attributed to specific characteristics of the datasets. For instance, the failure to fully segregate the protease dataset in Group 2 and all the datasets in Group 4 may be due to fine-grained functional differences within the datasets. Similarly, the challenge with the GPCR1 dataset in Group 5 may arise from the presence of transmembrane domains in both classes, which share a similar amino acid composition dominated by apolar residues.

These observations suggest that clustering methods are less effective in distinguishing fine-grained functional differences within protein families. However, the ability of the methods to differentiate Group 1 datasets, where each class was confined to a single glycoside hydrolase (GH) family, suggests that clustering is effective in distinguishing GH families despite functional similarities. This conclusion was further reinforced by similar results in Group 3, although the separation was only partial, potentially due to the presence of similar protein folds, a factor not observed in Group 1. Additionally, clustering methods are failing to segregate GH families in Group 4. Based on these results, clustering methods show promise as a tool for labeling unknown proteins, particularly for distinguishing GH families with fine-grained functional similarities.

For affinity propagation clustering (APC), we did not apply either intrinsic or extrinsic evaluation metrics. The intrinsic metric was not used since the optimal number of clusters *k* is determined by the algorithm itself, and the extrinsic metric was not applied because APC produced more clusters than expected. This resulted in a high number of false negatives and a lower FMI score. Nonetheless, the results obtained from APC were largely consistent with the trends observed in the combination of other algorithms.

### Current approaches for functional annotations

Supervised learning methods have also been used for protein function annotation using amino acid sequence as input. FAPM (Xiang et al., 2024) [12] is a contrastive multimodal model which combines a pretrained protein sequence model with a pretrained large language model. It generates GO functional terms, for example, UMP-CMP kinase, DNA binding, peptidase activity, ubiquitin protein ligase activity, pinoresinol reductase activity, transcription factor activity, electron transfer activity, and so on. GPSFun (Yuan et al., 2024) [13] is a geometry-aware model. It predicts 3D structures using a language model and a graph neural network. Using this, binding sites for ligands such as ATP, heme, metal ions, and other macromolecules such as DNA, RNA, and proteins are predicted. In addition, subcellular localization and solubility are also predicted. In both these cases, function prediction is rather coarse-grained. EnzBert (Buton et al., 2023) [14] is a transformer-based model that predicts functions upto 4th digit of the EC number. These three studies use large training datasets, labels from experimental and homology-based annotations, and employ complex protein language models and deep learning architectures, which are computationally intensive. In contrast to these approaches, (i) large data, (ii) labeled data, and/or (iii) heavy resources are not mandatory for unsupervised clustering as used in our study. Another key contrast is that data is manually curated by subject matter experts are used, leaving no scope for propagation of errors that have crept in in databases. Clearly, this approach requires intensive “human resources” and hence is not scalable. The two approaches viz., (i) scalable, supervised methods and (ii) unsuperived clustering methods working with small, well curated datasets (as present in our study) are complementary in their advantages and disadvantages.

Beside our study, there have also been other efforts to use unsupervised learning methods for functional annotations. For example, Littmann and coworkers [15] improved functional consistency of functional families in the CATH database using embeddings from ProtBERT and DBSCAN as the clustering algorithm. DPCfam [16] clustered 23 million UniRef50 sequences via density peak clustering from BLAST alignments. This recovered 81% of medium-to-large Pfam families and found 14,000 clusters with no existing Pfam annotation. DeepSeqProt [17] is an unsupervised deep-learning framework that can cluster proteins according to the GO-term annotations. DeepSeqProt results in fewer and larger clusters of proteins, which generally share functional similarity; these may belong to multiple protein families.

These studies primarily utilize complex, high-dimensional embedding. In contrast, our approach focuses on handcrafted features like pseudo amino acid composition (PAAC), allowing direct attribution of clustering outcomes to biologically meaningful properties. Our benchmarking of multiple clustering algorithms (e.g., k-means clustering, Gaussian mixture model, agglomerative clustering) reveals which algorithms are best suited for functional separation of proteins under various conditions. This aligns with the findings of Rodriguez et al. (2019) [18] where they systematically compared clustering algorithms on various types of synthetic datasets and highlighted that algorithms’ choice and parameter tuning significantly affect clustering outcomes.

### Application of the proposed approach on large protein datasets

How does the approach presented in the study work if the number of datapoints used as input for clustering is large? The GPCR dataset used in this study has about ≈3500 sequences, and by “large” we mean a dataset with *>*3500 sequences. As noted in the subsection titled Runtimes of clustering in Supporting information S1 File, the runtime is *<*13 seconds even when the sequences are ≈3500. From this, we infer that the runtime will not matter for datasets that are large in terms of size. However, what matters is the granularity of functional difference. This may be discerned by comparing the outcome for the protease (83 sequences) and GPCR2 (3558 sequences) datasets (Tables 4 and 5, in Supporting information S1 File).

The approach has similar applications as those for any other clustering approach and is independent of the size of the dataset. Some of the applications are given below.

i. Cluster assignment of new sequences: Once clusters are defined for a representative dataset, new sequences can be assigned using predictive parameter like predict()in scikit-learn for algorithms such as k-means and GMM. For clustering methods lacking predictive parameter (e.g., agglomerative or spectral), the dataset can be reclustered with new sequences included. This strategy also used in resources like Clusters of Orthologous Groups of proteins (COGs) [19] and NCBI’s ProtClustDB [20]. It is to be noted that “new sequences” here mean sequences without annotations.
ii. Pre-filtering for profile construction: Clustering can serve as a preprocessing step to group similar proteins, a prerequisite for building multiple sequence alignments and profile hidden Markov models (HMMs). For instance, protein clusters from resources like ProtClustDB [20] have been used to construct the NCBIfam-PRK HMM library, which supports large-scale functional annotation of prokaryotic genomes [21]. These clustered protein sets form the basis for generating profile HMMs, which are widely used for accurate and scalable annotation across microbial genomes.
iii. Feature quality assessment: Clustering also functions as a diagnostic tool for evaluating feature quality. If clustering on a given feature set yields high-quality groupings (e.g., as judged by SC or FMI), it suggests that the features capture relevant biological similarities. Thus, clustering provides a way to benchmark representations without relying on labeled data.
iv. Exploratory annotation of large-scale datasets: Clustering enables discovery of novel families and remote homologs in large, unlabeled datasets. For example, the Foldseek cluster approach [22], though structure-based, demonstrates how clustering can be scaled to millions of proteins to uncover novel structure-function relationships. While our study is sequence-based, the same principle applies: clustering provides a scalable and interpretable framework for organizing protein sequence space.

## Conclusion

This study aimed to assess the ability of clustering algorithms to segregate functionally distinct protein sequences using pseudo amino acid composition (PAAC) as the feature set.

Among the clustering algorithms tested, agglomerative performed slightly better than k-means and Gaussian mixture model (GMM) as assessed by the silhouette coefficient and Fowlkes-Mallows index. This could be due to the ability of agglomerative clustering to mimic phylogenetic relationships. However, k-means and GMM also showed a good capacity to reflect phylogeny to a large extent, and all three algorithms performed comparably well. While one may outperform the others in specific cases, we recommend using all three algorithms (k-means, GMM, and agglomerative) across a range of *k* values, choosing the optimal *k* based on the SC. For datasets containing proteins with known functions, the FMI can be used to select the best clusters and transfer annotations to unannotated proteins. However, if all data points are of unknown function, SC will be the sole metric for guiding cluster selection.

In the case of affinity propagation clustering (APC), while it performed well overall, its tendency to split homologous proteins into multiple clusters indicates higher sensitivity. For a more comprehensive analysis, it is advisable to use a combination of k-means, GMM, agglomerative, and APC, and compare the results for consistency. This approach provides a robust framework for the clustering and functional annotation of protein sequences, especially when dealing with unannotated datasets.

## Abbreviations

GH: glycoside hydrolases
FDX: ferredoxin
TM: transmembrane
PAAC: pseudo-amino acid composition
SC: silhouette coefficient
CD: Calinski-Harabasz
DB: Davies-Bouldin
GMM: Gaussian mixture model
APC: affinity propagation clustering
FMI: Fowlkes-Mallows index

## Acknowledgment

We thank Indian Institute of Technology Bombay for infrastructure and other facilities.

## Supporting information

**S1 File. Details on dataset and clustering analysis**. The file contain details of dataset creation, feature engineering, clustering algorithms, and values used for their parameters, assessing the suitability of different intrinsic and extrinsic metrics, and the reasons for not considering mean-shift, DBSCAN, OPTICS, and BIRCH algorithms, along with relevant data to support the main text.

