## Supplementary material for "How suitable are clustering methods for functional annotation of proteins?": S1 File

### Details related to datasets

#### Nomenclature

A protein sequence is treated as a 'datapoint' and a protein family is treated as a 'class' in the context of the clustering algorithm. Homologs are the proteins that evolved from the same ancestral protein. Among homologs, orthologs are those that retain molecular function, while paralogs diverge functionally; the extent of functional divergence varies [1]. A protein family is composed of a set of sequence or structural homologs. Proteins that perform the same molecular function are referred to as functional homologs. Functional homology can arise even in the absence of shared ancestry due to convergent evolution. In this research paper, "function" specifically refers to "molecular function", one of the three "aspects" used by Gene Ontology consortium to organize biological knowledge about proteins; the other two are the cellular component and the biological process. Molecular function covers reaction catalyzed (EC number), ligand/substrate specificity, etc. [2].

### Creation of datasets

We manually curated the datasets as described here. It was observed that the sequence lengths of proteins within a protein family show large variation. Therefore, outliers identified using a box-whisker plot were omitted. The following sequences were also omitted: (i) contain letters B, J, O, U, X, and Z, since these letters do not uniquely denote any of the 20 standard amino acids, (ii) have the term "fragment" or "partial" in their annotation, and (iii) have two or more EC numbers in their annotation. Additionally, within a protein family, only one copy of identical sequences or accession numbers was retained, while identical sequences or accession numbers were removed across protein families within the dataset. Finally, sequences with a length <30 were omitted because the default value of lambda was set to 30 for calculating the PAAC of sequences, resulting in a uniform 50-dimensional feature vector.

CAZy is a database of carbohydrate-active enzymes wherein protein sequences have been grouped into different families based on sequence similarity. The protein families in Xylanase, Chitinase, $\beta$-Glucosidase, Lysozyme, $\beta$-Galactosidase, and Lysozyme CGCh datasets were fetched from the CAZy database [3] using their corresponding EC numbers as queries. Additionally, protein names were used for the protein families in the Lysozyme CGCh dataset. For other families, protein names could not be used because of the multiple and ambiguous nomenclatures. These datasets were chosen due to their relatively large size. The GH2, GH3, and GH5 datasets were also taken from the CAZy database due to their large size and the presence of two functionally distinct protein families. Protein families in the Protease dataset were fetched from BRENDA [4] using EC numbers, and those in the Ferredoxin dataset from Swiss-Prot [5] using protein family names. $\alpha$-Lactalbumin sequences were collected from [6] and these constitute the Lysozyme CaLA dataset along with awC sequences.

The SNARE dataset is identical to the D128 dataset described by [7] and was obtained from their electronic supplementary data. The authors of this work downloaded SNARE sequences from UniProt using the filter GO:0005484, which corresponds to the molecular function "SNAP receptor activity," while non-SNARE sequences were fetched from the PDB database [8]. They further used CD-HIT [9] with a 25% sequence similarity threshold to obtain a non-redundant dataset. The SNARE family within the SNARE dataset consists of proteins that perform a generic receptor function but differ from each other in various aspects: localization, key functional residue, presence of transmembrane domains, coiled-coil domain, downstream pathway, etc. [10]; there is no discernible sequence identity with one another.

The GPCRs in GPCR1 and GPCR2 datasets were obtained from Swiss-Prot using the advanced search option. The "protein family" field under "Family and Domains" was queried with the keywords 'G protein-coupled receptor $n$ family' (where $n = 1,\ldots,5$), 'G protein-coupled receptor fz smo family', and 'G protein-coupled receptor T2R family', which correspond to IUPHAR classes A-F and Taste 2, respectively [11],[12]. The non-GPCRs were also obtained from Swiss-Prot using the NOT operator for the keywords given above. They were then split into two groups: (i) sequences with at least one transmembrane domain (non-GPCR(TM)) and (ii) sequences with no transmembrane domain (non-GPCR(no TM)). A 90% sequence identity cut-off was applied to each of the three protein families using CD-HIT to reduce redundancy and size. From these, 2000 entries were chosen randomly from each of the non-GPCR(TM) and non-GPCR(no TM) families in order to maintain class balance with the GPCR family in the GPCR1 and GPCR2 datasets, respectively. Details of the above mentioned 15 datasets and their associated protein families are shown in Table 1.

### Sequence similarity across protein families of a dataset

We performed an all-against-all sequence comparison using BLASTp. Two proteins were considered to be similar (or homologous) if the percentage sequence identity (PSI) $\geq$30% [17] and the alignment covers at least 70% (empirically) of one of the sequences. This was used to infer whether protein families shared sequence similarity or not (data not shown).

**Table 1. Groups, datasets, and protein families with associated functions, structures, and number of sequences.**

| Group | Dataset | Protein family | EC # | Fold | $N_{seq}$ |
|---|---|---|---|---|---|
| Group 1 | Lysozyme CGCh | Lysozyme C | 3.2.1.17 | lysozyme fold | 52 |
| | | Lysozyme G | 3.2.1.17 | lysozyme fold | 17 |
| | | Lysozyme Ch | 3.2.1.17 | $(\beta/\alpha)_5(\beta)_3$ Tim-like | 54 |
| | Xylanase | GH10 | 3.2.1.8 | $(\beta/\alpha)_8$ barrel | 352 |
| | | GH11 | 3.2.1.8 | $\beta$-jelly roll | 347 |
| | Chitinase | GH18 | 3.2.1.14 | $(\beta/\alpha)_8$ barrel | 462 |
| | | GH19 | 3.2.1.14 | lysozyme fold | 194 |
| Group 2 | Lysozyme CaLA | Lysozyme C | 3.2.1.17 | lysozyme fold | 52 |
| | | $\alpha$-Lactalbumin | None | lysozyme fold [13] | 22 |
| | Protease | Trypsin | 3.4.21.4 | chymotrypsin/ trypsin fold [14] | 67 |
| | | Chymotrypsin | 3.4.21.1 | chymotrypsin/ trypsin fold | 16 |
| | Ferredoxin | FDX1 | None | ferredoxin fold [15] | 9 |
| | | FDX2 | None | ferredoxin fold | 7 |
| Group 3 | $\beta$-Glucosidase | GH1 | 3.2.1.21 | $(\beta/\alpha)_8$ barrel | 348 |
| | | GH3 | 3.2.1.21 | $(\beta/\alpha)_8$ barrel | 204 |
| | Lysozyme | GH22 | 3.2.1.17 | lysozyme fold | 135 |
| | | GH23 | 3.2.1.17 | lysozyme fold | 47 |
| | | GH24 | 3.2.1.17 | lysozyme fold | 31 |
| | $\beta$-Galactosidase | GH2 | 3.2.1.23 | $(\beta/\alpha)_8$ barrel | 117 |
| | | GH35 | 3.2.1.23 | $(\beta/\alpha)_8$ barrel | 295 |
| | | GH42 | 3.2.1.23 | $(\beta/\alpha)_8$ barrel | 83 |
| Group 4 | GH2 | $\beta$-Galactosidase | 3.2.1.23 | $(\beta/\alpha)_8$ barrel | 117 |
| | | $\beta$-Glucuronidase | 3.2.1.31 | $(\beta/\alpha)_8$ barrel | 85 |
| | GH3 | $\beta$-Glucosidase | 3.2.1.21 | $(\beta/\alpha)_8$ barrel | 204 |
| | | 1,4-$\beta$-Xylosidase | 3.2.1.37 | $(\beta/\alpha)_8$ barrel | 99 |
| | GH5 | Cellulase | 3.2.1.4 | $(\beta/\alpha)_8$ barrel | 342 |
| | | Endo-$\beta$-Mannanase | 3.2.1.78 | $(\beta/\alpha)_8$ barrel | 127 |
| Group 5 | SNARE | SNARE | None | No specific fold | 57 |
| | | non-SNARE | None | No specific fold | 61 |
| | GPCR1 | GPCR | None | 7TM helix fold [16] | 1697 |
| | | non-GPCR(TM) | None | No specific fold | 1907 |
| | GPCR2 | GPCR | None | 7TM helix fold | 1697 |
| | | non-GPCR(no TM) | None | No specific fold | 1861 |

The table lists all five groups, the datasets within each group, and the protein families in each dataset. For each protein family, the function (EC number (EC#)), structure (Fold), and number of sequences ($N_{seq}$) are shown. The 5 groups comprise a total of 15 datasets (3 datasets per group), with each dataset containing two or three protein families.

### Feature engineering

### Pseudo amino acid composition (PAAC)

Sequences of proteins must be transformed into feature vectors of fixed dimensions for clustering algorithms to process them. Pseudo amino acid composition is a widely used and robust feature extraction method that captures both amino acid composition and sequence-order correlation [18]. In our study, we used the default values from the original paper [19] for the weightage factor (0.05) and lambda (30, as chosen intuitively); the physicochemical properties used are the same as those suggested, i.e., hydrophobicity, hydrophilicity, and side chain mass. As a result, each feature vector has 50 dimensions, where the first 20 correspond to amino acid composition, and the next 30 correspond to sequence-order correlation. We used in-house scripts for all data processing steps except where mentioned to the contrary.

### Details related to clustering algorithms

#### Algorithms

Clustering is a type of unsupervised learning [20]. We have assessed nine clustering algorithms provided in the scikit-learn platform [21]. The number of clusters $k$ to be formed by input data is user-defined for k-means clustering, spectral clustering, agglomerative clustering, and Gaussian mixture model (GMM). On the other hand, the number of clusters is determined by the algorithm itself in the case of DBSCAN clustering, OPTICS clustering, affinity propagation clustering (APC), and mean-shift clustering. In the case of BIRCH clustering, `n_clusters` was set to `X`, where `X` represents the number of classes in the dataset. BIRCH first creates a clustering feature tree (CFT) from input data and groups datapoints into subclusters. If the number of subclusters is greater than the user-defined $k$, then BIRCH uses a global clustering algorithm to reduce the number of subclusters to $k$. And if the number of subclusters is less than or equal to the user-defined $k$, then BIRCH retains the current subclusters as the final clusters [22]. All algorithms were used with their default values, except for two cases: (i) in spectral clustering, the default value of `affinity` parameter `rbf` was changed to `nearest_neighbors` to ensure scale invariance, (ii) in APC, the value for the `preference` parameter was changed for reasons given below.

The performance of APC is primarily dictated by the parameter `preference`. Choosing a `higher` value can result in more number of datapoints to be cluster centers (the so-called exemplars). Alternatively, one can assign `preference` values for each of the datapoints, and those with higher values are likely to be exemplars. By default, the `preference` is set to the median of the similarity matrix [23]. The number of clusters formed for various datasets ranged between 4 and 249. These numbers are very large when one considers the way datasets are curated; for example, at most 2 or 3 clusters were expected for the lysozyme CGCh dataset. Using the minimum value in the similarity matrix is an option, and this is expected to reduce the number of clusters. Hence, the datasets were clustered by setting `preference` to the minimum value. As expected, fewer clusters are obtained for all datasets.

### Assessment of the quality of clustering

The quality of clustering was assessed using (i) intrinsic metrics and (ii) database-assigned labels [24].

#### Assessment using intrinsic metrics

Different intrinsic metrics have been used by various research groups. In the present study, three intrinsic evaluation metrics available in the scikit-learn [21] platform were computed. These are the silhouette coefficient (SC) [25], Calinski-Harabasz (CH) index [26], and Davies-Bouldin (DB) index [27]. These three metrics were part of a group of 6 metrics (out of 30) that were found to perform satisfactorily against 720 synthetic and 20 real datasets clustering using k-means, ward, and average-linkage algorithms [28].

The SC is calculated by comparing the mean distance between a sample and all other points within the same cluster to the mean distance between the sample and points in the nearest neighboring cluster. The SC ranges from -1 to +1, where -1 indicates incorrect clustering, +1 indicates highly dense clustering, and values around zero suggest overlapping clusters [25]. The DB index compares the distance between clusters with the size of the clusters. Zero is the lowest possible score, which indicates a better partition. However, there is no defined upper bound for the DB index [27]. The CH index is the ratio of the sum of between-clusters dispersion and the sum of within-cluster dispersion for all clusters (here the dispersion is defined as the sum of squared distances) [26].

#### Assessment using protein sequence database-assigned labels

Even though clustering is an unsupervised learning method, database-assigned labels have also been used to assess the quality of clustering by means of contingency matrices and the Fowlkes-Mallows index (FMI) [29]. It may be noted that database-assigned labels are labels that have not been part of the input for clustering. For preparing contingency matrices, sequence database annotations of the input datapoints (i.e., protein sequences) were used as labels for comparison with the cluster labels assigned by clustering algorithms.

Extrinsic evaluation metrics are less commonly used than intrinsic metrics, mainly due to the requirement of database-assigned labels [28]. In this study, we considered four extrinsic metrics: FMI, F1, MCC, and AMI.

To check for redundancy among these metrics, we computed the correlation coefficient between every pair of metrics on all datasets and among all algorithms with $k$= 2-10. The correlation coefficients were found to be 0.99 between FMI and F1, 0.95 between AMI and MCC, and 0.77 between FMI and AMI. Hence, we only considered FMI for further analysis.

FMI [29] is the geometric mean of pairwise precision and recall. It is calculated by considering two datapoints at a time. This is illustrated in Table 2 where circles represent datapoints and their colors represent assigned labels. The class is calculated for all pairs of datapoints based on database-assigned labels and cluster labels. The cluster(s) to which these datapoints belong and the labels assigned from the database are compared. FMI is 1 when every pair of datapoints that belong to the same cluster has the same labels. FMI is 0 when every pair belonging to the same cluster has non-identical labels.

$$FMI = \frac{TP}{\sqrt{(TP+FP)(TP+FN)}} \quad (1)$$

**Table 2. Calculation of TP, TN, FP, and FN outcomes for unsupervised learning.**

| Outcome | Database-assigned labels | Cluster labels |
|---|---|---|
| True Positive | 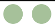 |  |
| True Negative | 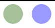 |  |
| False Negative | 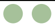 |  |
| False Positive | 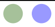 |  |

The table illustrates how True Positives (TP), True Negatives (TN), False Negatives (FN), and False Positives (FP) are determined for pairs of datapoints in an unsupervised learning context. Outcomes are computed by comparing identical or non-identical database-assigned labels with identical or non-identical cluster labels. For example, a pair with identical database-assigned labels and identical cluster labels is considered a TP. In the table, datapoints are represented as circles, and colors indicate their labels.

### Suitability of intrinsic metrics and possible thresholds

As mentioned in the previous section, the silhouette coefficient (SC), Calinski-Harabasz (CH) index, and Davies-Bouldin (DB) index were used to analyze clustering output. For a given combination of (dataset + algorithm), each of the three metrics was plotted as a function of $k$.

By definition, SC is bounded between -1 for incorrect clustering and +1 for highly dense clustering [25]. Scores around zero indicate overlapping clusters. SC was computed for all combinations of datasets, clustering algorithms (k-means, GMM, agglomerative, and spectral clustering), and $k$ (varied from 2 to 10). It was observed that SC is not close to 1 for any of the clusters. It is not clear if this is because (i) all the clusters are of poor quality, (ii) if this is the best separation one can get for the type of datasets + features we have used, or (iii) extent of sequence similarity between protein families vis-a-vis within protein families. These were resolved as follows:

(i) We clustered a set of datapoints that consists of two completely unrelated protein families, i.e., SNARE and trypsin. It is seen that SC is between 0.31 and 0.32.

(ii) We assigned database-assigned labels to every datapoint and treated these as the cluster labels for computing the SC value. Taking the Xylanase dataset as an example, the computed SC value was 0.35, which closely matches the SC values obtained using cluster labels from the four algorithms, which ranged from 0.35 to 0.36 for $k = 2$. A similar pattern was observed for other datasets also.

(iii)We computed the inter/intra similarity ratio for each dataset to assess whether clustering quality (measured by the SC) is related to sequence similarity. This ratio is defined as the mean sequence identity between protein families divided by the mean sequence identity within protein families in the dataset. A low inter/intra ratio indicates that families are well separated in sequence space (i.e., low inter-family similarity and high intra-family similarity). In such cases, we generally expect higher SC values, leading to a negative correlation between the inter/intra ratio and SC.

When plotting the SC values against the inter/intra similarity ratios across datasets (Fig 1), we did observe an overall negative trend, although several exceptions were noted. This suggests that sequence similarity alone does not fully explain clustering performance. Notably, none of the datasets had an inter/intra ratio below 0.6, indicating that even in the best cases, protein families were not entirely dissimilar at the sequence level.

From these, we infer that SC is unlikely to reach the highest value (1.00) irrespective of the value of $k$ and the clustering algorithm for the type of data and feature engineering we are using in this study. In view of this, for a given algorithm, we chose the $k$ value for which SC is highest and positive as optimal. This $k$ value happens to be the same if we were to use the lowest DB index instead of the highest SC, except in 8 out of 15 cases. The choice of SC over the DB index is because the DB index does not have an upper limit.

The CH index decreases monotonically as $k$ increases, with a few exceptions, as shown in Fig 2. Thus,

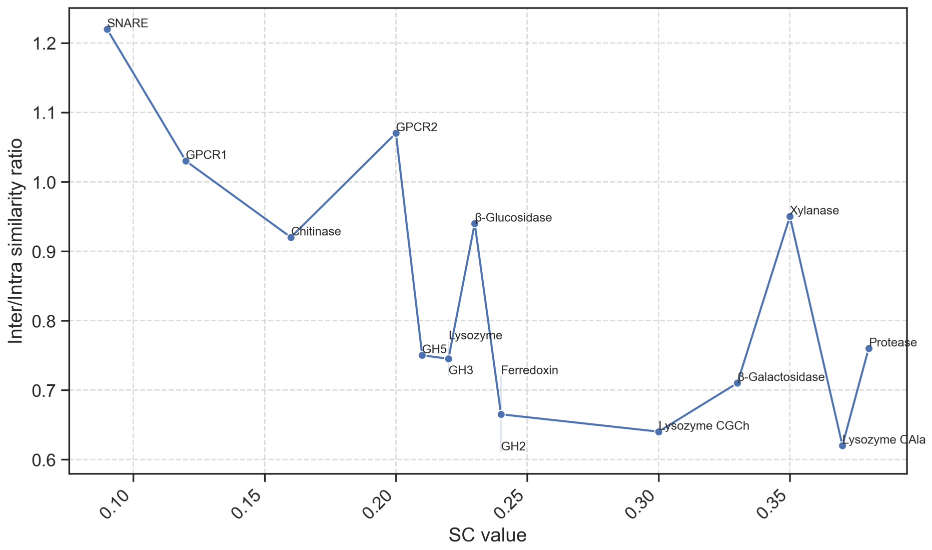


**Fig 1. Comparison of silhouette coefficients and inter/intra similarity ratios for 15 datasets.** The line plot compares silhouette coefficients (SC) and inter/intra similarity ratios for 15 datasets. As a representative case, SC values were obtained using agglomerative clustering, with the number of clusters ($k$) set to the number of protein families in each dataset. Points in the graph are labeled with the dataset name corresponding to their SC value and inter/intra similarity ratio.

$k = 2$ happens to have the best score irrespective of the input dataset and the clustering algorithm. Also, by definition, there is no upper bound for the CH index. The highest CH index for the various datasets (among all algorithms) ranged from 6.8 (Ferredoxin, $k = 10$, agglomerative) to 1001.3 (GPCR2, $k = 2$, k-means) as shown in Fig 3. It is not clear how this index can be used to ascertain the quality of clustering when one is working with multiple input datasets and clustering algorithms.

Overall, we infer that the CH index and DB index are not suitable in the current study. SC was used without reference to the absolute value as long as it is positive.

### Analysis of the number and size of clusters formed by mean-shift, DBSCAN, OPTICS, and BIRCH

Mean-shift, DBSCAN, OPTICS, and BIRCH determine the number of clusters to be formed based on input data. The number of clusters formed is shown in Table 3. BIRCH gave a single cluster for all the datasets. Mean-shift groups most of the datapoints in a single cluster, even when the number of clusters formed is >1. DBSCAN and OPTICS identify most of the datapoints as noise points irrespective of the number of datapoints. This output cannot be reconciled considering domain knowledge as explained in detail below.

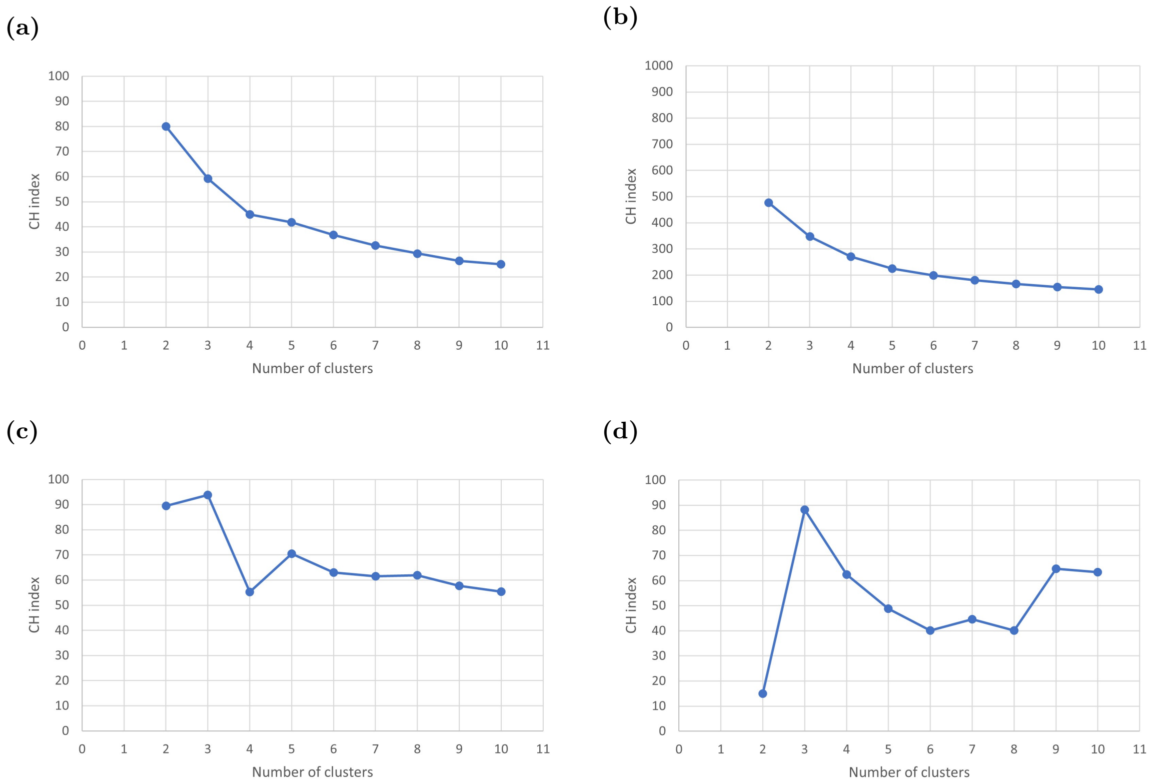


**Fig 2. Variation of Calinski–Harabasz index with number of clusters in representative datasets and algorithms.** The line plots show how the Calinski–Harabasz (CH) index changes as the number of clusters ($k$) increases from 2 to 10 across four representative dataset-algorithm combinations. The CH index decreases monotonically with increasing $k$ in most cases, as seen in (a) k-means clustering of the Lysozyme CGCh dataset and (b) agglomerative clustering of the Xylanase dataset. Exceptions include (c) spectral clustering of the Chitinase dataset and (d) spectral clustering of the $\beta$-Glucosidase dataset. Data for other datasets and algorithms are not shown.

The performance of mean-shift is primarily dictated by the parameter `bandwidth`, which determines the size of the window in which datapoints are considered for calculating the mean. By default, mean-shift uses an estimator to determine the value of the `bandwidth`. It appears that the algorithm has estimated a high `bandwidth` value, resulting in larger neighborhoods leading to a single cluster.

The performance of DBSCAN is primarily dictated by the parameters `eps` and `min_samples`. The former sets the maximum distance between two samples for one to be considered as in the neighborhood of the other. The latter determines the number of samples in a neighborhood for a point to be considered as a core point. There is no basis on which values for these parameters can be chosen for the datasets considered in this study. Hence, default values, i.e., 0.5 and 5, respectively, were chosen. With these values, while a single cluster was obtained for some datasets, 6-28 clusters were obtained for the remaining datasets.

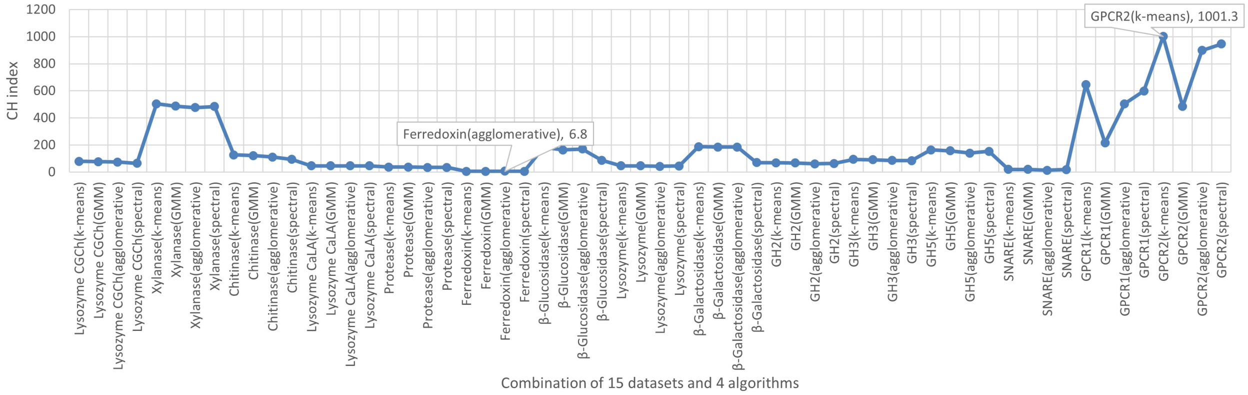


**Fig 3. Highest Calinski–Harabasz index across datasets and clustering algorithm.** The line plot shows the highest Calinski–Harabasz (CH) index for each combination of 15 datasets and 4 clustering algorithms. Values range from 6.8 for the Ferredoxin dataset ($k$=10, agglomerative clustering) to 1001.3 for the GPCR2 dataset ($k$=2, k-means clustering).

**Table 3. Number of clusters and datapoints in the largest cluster for three clustering algorithms across datasets.**

| Group | Dataset | # seq. | mean-shift | | DBSCAN | | OPTICS | |
|---|---|---|---|---|---|---|---|---|
| | | | $k$ | $N_{largest}$ | $k$ | Noise | $k$ | Noise |
| Group 1 | Lysozyme CGCh | 123 | 3 | 121 | 1 | 123 | 6 | 72 |
| | Xylanase | 699 | 5 | 365 & 331$^{\sigma}$ | 14 | 610 | 38 | 418 |
| | Chitinase | 656 | 5 | 633 | 24 | 504 | 48 | 301 |
| Group 2 | Lysozyme CaLA | 74 | 5 | 46 & 24$^{\sigma}$ | 1 | 74 | 5 | 50 |
| | Protease | 83 | 1 | 83 | 1 | 83 | 5 | 51 |
| | Ferredoxin | 16 | 4 | 12 | 1 | 16 | 1 | 16 |
| Group 3 | $\beta$-Glucosidase | 552 | 1 | 552 | 19 | 425 | 32 | 322 |
| | Lyszoyme | 213 | 1 | 213 | 8 | 170 | 15 | 77 |
| | $\beta$-Galactosidase | 495 | 5 | 418 | 28 | 250 | 49 | 143 |
| Group 4 | GH2 | 202 | 1 | 202 | 7 | 147 | 12 | 108 |
| | GH3 | 303 | 1 | 303 | 10 | 243 | 18 | 182 |
| | GH5 | 469 | 2 | 467 | 6 | 437 | 20 | 345 |
| Group 5 | SNARE | 118 | 5 | 113 | 1 | 118 | 1 | 118 |
| | GPCR1 | 3604 | 84 | 3351 | 1 | 3604 | 32 | 3376 |
| | GPCR2 | 3558 | 92 | 3278 | 1 | 3558 | 29 | 3353 |

The table shows the number of clusters ($k$) and the total number of datapoints in the largest cluster ($N_{largest}$) for mean-shift, DBSCAN, and OPTICS applied to each of the 15 datasets in 5 groups. For DBSCAN and OPTICS, the largest clusters consisted of noise points (Noise).

$^{\sigma}$ In this case, we showcased two clusters with maximum datapoints.

The performance of OPTICS is primarily determined by the parameters `max_eps` and `min_samples`. Being a variant of DBSCAN, `max_eps` has the same meaning as `eps`. The default value for this parameter is $\infty$. The use of default values led to a single cluster for two datasets (Ferredoxin and SNARE) and 5–49 clusters for other datasets.

The performance of BIRCH is primarily determined by the parameters `threshold` and `branching_factor`. The former parameter (default value = 0.5) defines the maximum radius within which a new sample can merge with the closest existing subcluster. The latter parameter (default value = 50) determines the maximum number of subclusters that can branch from a parent cluster in the hierarchical structure. A single cluster was obtained for all 15 datasets, suggesting that the default value for `threshold` is so high that even though `branching_factor` is set to 50, only a single cluster is formed.

It can be inferred that the default values for the various parameters of these four algorithms are not optimal for the feature engineering that we have chosen. Given the nature of the input considered in the present study, i.e., unlabeled datapoints whose relative spread in the 50-D space is not known a priori, there is no straightforward way to determine optimal values for the various parameters. Limited efforts to fine-tune the values were infructuous because values that may seem optimal for one dataset are not necessarily optimal for others. Hence, results from these four algorithms were not considered to arrive at the conclusions.

Overall, it is concluded that the output from these four algorithms will not be considered for further analysis.

#### Patterns of cluster formation by k-means, GMM, agglomerative, and spectral clustering

All 15 datasets were used as input separately for each of these four algorithms and by varying $k$ from 2 to 10. The results were first analyzed by comparing the membership of datapoints in different clusters for two consecutive values of $k$ in Fig 4. It was observed that with each increment of $k$, agglomerative splits one, and only one, cluster into two. This is to be expected since agglomerative is a hierarchical method and, by design, iteratively clusters datapoints based on *mutual similarity* in the feature space. The process of clustering is depicted as a dendrogram and mimics a phylogenetic tree. In the present study, protein families that constitute datasets are indeed phylogenetically related. Hence, splitting of clusters with increments in $k$ can be thought of as depicting evolutionary divergence.

The results from the other three algorithms show that the pattern of splitting is somewhat different, i.e., when $k$ increases by 1, some of the newly formed clusters have datapoints from $\geq$2 clusters. This shows that the association of any two datapoints that are part of the same cluster is not guaranteed to remain when $k$ changes. However, in most cases (i.e., for a given combination of dataset + algorithm + $k$), only $\leq$10% of datapoints show this behavior; in 4-5 cases, we found up to 30% of datapoints change cluster

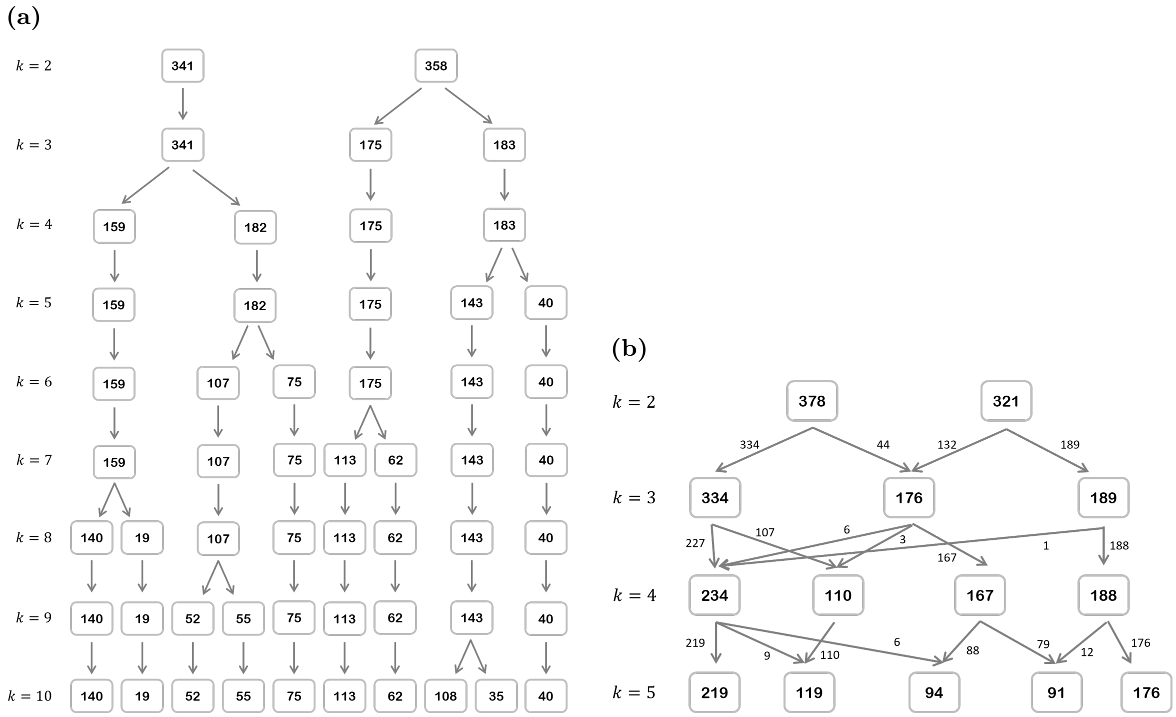


**Fig 4. Cluster splitting patterns with increasing number of clusters for clustering algorithms.** The figure illustrates how clusters split as the number of clusters ($k$) increases from 2 to 10 for two clustering algorithms: (a) agglomerative clustering and (b) k-means clustering on the Xylanase dataset.

membership, and this often happened for higher values of $k$. These observations suggest that the pattern of cluster formation mimics phylogeny, especially for lower values of $k$.

### Reasons for using SC and FMI to assess clustering

We find that the outcome of clustering methods is critically dependent on four factors: (i) type, nature, and size of data, (ii) feature vector, (iii) clustering algorithms, and (iv) parameters used by algorithms. For a given task, the type of data is fixed: in this case, the data are protein sequences. In the present study, the nature of the data refers to sequence variability and the relationship between sequence and function. The latter is quite subtle and is not fully understood, especially when the focus is on fine-grained functional differences (e.g., substrate specificity). In an ideal situation, unsupervised learning algorithms ought to be assessed by intrinsic metrics, but these can be biased by the spread of data in the feature vector space. Thus, there is no universally applicable metric, i.e., a metric that is suitable for all combinations of types of data, feature vector, and clustering algorithm. Hence, extrinsic metrics are often used, and these also have limitations. Consequently, there is no way to find the most appropriate combination of feature vector, algorithm, and parameter values for a given dataset. To address these challenges, in the present study, we opt for a feedback-driven iterative approach to find the optimal combination, and we used both intrinsic (silhouette coefficient) and extrinsic (Fowlkes-Mallows index) metrics.

**Table 4. Contingency matrices of clustering results with the highest Fowlkes–Mallows index.**

| Group | Protein family | Cluster label | | | | | | | | | | Total |
|---|---|---|---|---|---|---|---|---|---|---|---|---|
| | | **0** | **1** | **2** | **3** | **4** | **5** | **6** | **7** | **8** | **9** | |
| 1 | (a) Agglomerative clustering of the Lysozyme CGCh dataset (FMI=0.88) | | | | | | | | | | | |
| | **Lysozyme C** | 52 | 0 | NA (only 2 clusters) | | | | | | | | 52 |
| | **Lysozyme G** | 17 | 0 | | | | | | | | | 17 |
| | **Lysozyme Ch** | 0 | 54 | | | | | | | | | 54 |
| | **Total** | 69 | 54 | | | | | | | | | 123 |
| | (b) Agglomerative clustering of the Xylanase dataset (FMI=0.98) | | | | | | | | | | | |
| | **GH10** | 351 | 1 | NA (only 2 clusters) | | | | | | | | 352 |
| | **GH11** | 7 | 340 | | | | | | | | | 347 |
| | **Total** | 358 | 341 | | | | | | | | | 699 |
| | (c) Spectral clustering of the Chitinase dataset (FMI=0.85) | | | | | | | | | | | |
| | **GH18** | 439 | 23 | NA(only 2 clusters) | | | | | | | | 462 |
| | **GH19** | 43 | 151 | | | | | | | | | 194 |
| | **Total** | 482 | 174 | | | | | | | | | 656 |
| 2 | (d) K-means, GMM, agglomerative, and spectral clustering of the Lysozyme CaLA dataset (FMI=1.00) | | | | | | | | | | | |
| | **$\alpha$-Lactalbumin** | 22 | 0 | NA (only 2 clusters) | | | | | | | | 22 |
| | **Lysozyme C** | 0 | 52 | | | | | | | | | 52 |
| | **Total** | 22 | 52 | | | | | | | | | 74 |
| | (e) Agglomerative and spectral clustering of the Protease dataset (FMI=0.70) | | | | | | | | | | | |
| | **Trypsin** | 56 | 11 | NA (only 2 clusters) | | | | | | | | 67 |
| | **Chymotrypsin** | 16 | 0 | | | | | | | | | 16 |
| | **Total** | 72 | 11 | | | | | | | | | 83 |
| | (f) Spectral clustering of the Ferredoxin dataset (FMI=0.75) | | | | | | | | | | | |
| | **FDX1** | 1 | 8 | NA (only 2 clusters) | | | | | | | | 9 |
| | **FDX2** | 6 | 1 | | | | | | | | | 7 |
| | **Total** | 7 | 9 | | | | | | | | | 16 |
| 3 | (g) Agglomerative clustering of the $\beta$-Glucosidase dataset (FMI=0.78) | | | | | | | | | | | |
| | **GH1** | 287 | 1 | 60 | NA (only 3 clusters) | | | | | | | 348 |
| | **GH3** | 21 | 153 | 30 | | | | | | | | 204 |
| | **Total** | 308 | 154 | 90 | | | | | | | | 552 |
| | (h) K-means and GMM of the Lysozyme dataset (FMI=0.43) | | | | | | | | | | | |
| | **GH22** | 0 | 4 | 40 | 17 | 17 | 10 | 0 | 26 | 21 | 0 | 135 |
| | **GH23** | 25 | 0 | 0 | 0 | 0 | 0 | 7 | 2 | 0 | 13 | 47 |
| | **GH24** | 6 | 0 | 0 | 0 | 3 | 0 | 22 | 0 | 0 | 0 | 31 |
| | **Total** | 31 | 4 | 40 | 17 | 20 | 10 | 29 | 28 | 21 | 13 | 213 |
| | (i) GMM of the $\beta$-Galactosidase dataset (FMI=0.86) | | | | | | | | | | | |
| | **GH2** | 74 | 0 | 43 | NA (only 3 clusters) | | | | | | | 117 |
| | **GH35** | 5 | 283 | 7 | | | | | | | | 295 |
| | **GH42** | 39 | 0 | 44 | | | | | | | | 83 |
| | **Total** | 118 | 283 | 94 | | | | | | | | 495 |

For each dataset, among four optimal $k$ values (one per algorithm), the clustering result with the highest Fowlkes–Mallows index (FMI) was chosen as the final result. This table shows the contingency matrices for all datasets present in Groups 1-3. A visual representation of the corresponding datapoint distributions in reduced-dimensional space is shown in Figs 7-9.

**Table 5. Contingency matrices of clustering results with the highest Fowlkes–Mallows index.**

| Group | Protein family | Cluster label | | | | | | | | | | Total |
|---|---|---|---|---|---|---|---|---|---|---|---|---|
| | | **0** | **1** | **2** | **3** | **4** | **5** | **6** | **7** | **8** | **9** | |
| 4 | (j) K-means, GMM, and agglomerative clustering of the GH2 dataset (FMI=0.37) | | | | | | | | | | | |
| | **$\beta$-Glucuronidase** | 27 | 29 | 5 | 0 | 4 | 2 | 1 | 4 | 5 | 8 | 85 |
| | **$\beta$-Galactosidase** | 0 | 4 | 15 | 24 | 20 | 20 | 14 | 12 | 6 | 2 | 117 |
| | **Total** | 27 | 33 | 20 | 24 | 24 | 22 | 15 | 16 | 11 | 10 | 202 |
| | (k) Spectral clustering of the GH3 dataset (FMI=0.62) | | | | | | | | | | | |
| | **$\beta$-Glucosidase** | 163 | 41 | NA (only 2 clusters) | | | | | | | | 204 |
| | **1,4-$\beta$-Xylosidase** | 89 | 10 | | | | | | | | | 99 |
| | **Total** | 252 | 51 | | | | | | | | | 303 |
| | (l) Spectral clustering of the GH5 dataset (FMI=0.58) | | | | | | | | | | | |
| | **Cellulase** | 114 | 228 | NA (only 2 clusters) | | | | | | | | 342 |
| | **Endo-$\beta$-Mannanase** | 63 | 64 | | | | | | | | | 127 |
| | **Total** | 177 | 292 | | | | | | | | | 469 |
| 5 | (m) K-means and GMM of the SNARE dataset (FMI=0.86) | | | | | | | | | | | |
| | **SNARE** | 1 | 56 | NA (only 2 clusters) | | | | | | | | 57 |
| | **non-SNARE** | 53 | 8 | | | | | | | | | 61 |
| | **Total** | 54 | 64 | | | | | | | | | 118 |
| | (n) GMM of the GPCR1 (FMI=0.66) | | | | | | | | | | | |
| | **GPCR** | 1692 | 5 | NA (only 2 clusters) | | | | | | | | 1697 |
| | **non-GPCR(TM)** | 1635 | 272 | | | | | | | | | 1907 |
| | **Total** | 3327 | 277 | | | | | | | | | 3604 |
| | (o) Agglomerative clustering of the GPCR2 dataset (FMI=0.94) | | | | | | | | | | | |
| | **GPCR** | 1650 | 47 | NA (only 2 clusters) | | | | | | | | 1697 |
| | **non-GPCR(no TM)** | 55 | 1806 | | | | | | | | | 1861 |
| | **Total** | 1705 | 1853 | | | | | | | | | 3558 |

For each dataset, among four optimal $k$ values (one per algorithm), the clustering result with the highest Fowlkes–Mallows index (FMI) was chosen as the final result. This table shows the contingency matrices for all datasets present in Groups 4 and 5. A visual representation of the corresponding datapoint distributions in reduced-dimensional space is shown in Figs 10 and 11.

**Table 6. Contingency matrices of affinity propagation clustering results.**

| Group | Protein family | Cluster label | | | | | | | | | | | | Total |
|---|---|---|---|---|---|---|---|---|---|---|---|---|---|---|
| | | **0** | **1** | **2** | **3** | **4** | **5** | **6** | **7** | **8** | **9** | **10** | **11** | |
| 1 | (a) APC of the Lysozyme CGCh dataset (SC=0.30) | | | | | | | | | | | | | |
| | **Lysozyme C** | 17 | 33 | 0 | 2 | NA (only 4 clusters) | | | | | | | | 52 |
| | **Lysozyme G** | 0 | 1 | 16 | 0 | | | | | | | | | 17 |
| | **Lysozyme Ch** | 0 | 0 | 3 | 51 | | | | | | | | | 54 |
| | **Total** | 17 | 34 | 19 | 53 | | | | | | | | | 123 |
| | (b) APC of the Xylanase dataset (SC=0.17) | | | | | | | | | | | | | |
| | **GH10** | 60 | 138 | 109 | 41 | 0 | 1 | 0 | 3 | 0 | NA (only 9 clusters) | | | 352 |
| | **GH11** | 0 | 0 | 3 | 0 | 80 | 59 | 92 | 45 | 68 | | | | 347 |
| | **Total** | 60 | 138 | 112 | 41 | 80 | 60 | 92 | 48 | 68 | | | | 699 |
| | (c) APC of the Chitinase dataset (SC=0.15) | | | | | | | | | | | | | |
| | **GH18** | 85 | 53 | 15 | 77 | 45 | 77 | 40 | 69 | 0 | 1 | NA (only 10 clusters) | | 462 |
| | **GH19** | 6 | 10 | 0 | 9 | 2 | 0 | 0 | 0 | 94 | 73 | | | 194 |
| | **Total** | 91 | 63 | 15 | 86 | 47 | 77 | 40 | 69 | 94 | 74 | | | 656 |
| 2 | (d) APC of the Lysozyme CaLA dataset (SC=0.29) | | | | | | | | | | | | | |
| | **Lysozyme C** | 8 | 11 | 33 | 0 | NA (only 4 clusters) | | | | | | | | 22 |
| | **$\alpha$-Lactalbumin** | 1 | 0 | 0 | 21 | | | | | | | | | 74 |
| | **Total** | 9 | 11 | 33 | 21 | | | | | | | | | 96 |
| | (e) APC of the Protease dataset (SC=0.23) | | | | | | | | | | | | | |
| | **Trypsin** | 14 | 5 | 20 | 28 | NA (only 4 clusters) | | | | | | | | 67 |
| | **Chymotrypsin** | 0 | 0 | 10 | 6 | | | | | | | | | 16 |
| | **Total** | 14 | 5 | 30 | 34 | | | | | | | | | 83 |
| | (f) APC of the Ferredoxin dataset (SC=0.20) | | | | | | | | | | | | | |
| | **FDX1** | 9 | 0 | NA (only 2 clusters) | | | | | | | | | | 9 |
| | **FDX2** | 2 | 5 | | | | | | | | | | | 7 |
| | **Total** | 11 | 5 | | | | | | | | | | | 16 |
| 3 | (g) APC of the $\beta$-Glucosidase dataset (SC=0.19) | | | | | | | | | | | | | |
| | **GH1** | 61 | 58 | 53 | 73 | 17 | 81 | 0 | 1 | 3 | 1 | NA (only 10 clusters) | | 348 |
| | **GH3** | 0 | 0 | 0 | 0 | 0 | 0 | 61 | 28 | 35 | 80 | | | 204 |
| | **Total** | 61 | 58 | 53 | 73 | 17 | 81 | 61 | 29 | 38 | 81 | | | 552 |
| | (h) APC of the Lysozyme dataset (SC=0.19) | | | | | | | | | | | | | |
| | **GH22** | 19 | 36 | 26 | 42 | 12 | 0 | NA (only 6 clusters) | | | | | | 135 |
| | **GH24** | 0 | 0 | 0 | 1 | 0 | 30 | | | | | | | 31 |
| | **GH23** | 0 | 0 | 0 | 0 | 2 | 45 | | | | | | | 47 |
| | **Total** | 19 | 36 | 26 | 43 | 14 | 75 | | | | | | | 213 |
| | (i) APC of the $\beta$-Galactosidase dataset (SC=0.25) | | | | | | | | | | | | | |
| | **GH2** | APC failed to converge with default parameters | | | | | | | | | | | | 117 |
| | **GH42** | | | | | | | | | | | | | 83 |
| | **GH35** | | | | | | | | | | | | | 295 |
| | **Total** | | | | | | | | | | | | | 495 |

This table presents the contingency matrices of affinity propagation clustering (APC) for all the datasets present in Groups 1-3. A visual representation of the corresponding datapoint distributions in reduced-dimensional space is shown in Figs 12-14.

**Table 7. Contingency matrices of affinity propagation clustering results.**

| Group | Protein family | Cluster label | | | | | | | | | | | | Total |
|---|---|---|---|---|---|---|---|---|---|---|---|---|---|---|
| | | **0** | **1** | **2** | **3** | **4** | **5** | **6** | **7** | **8** | **9** | **10** | **11** | |
| 4 | (j) APC of the GH2 dataset (SC=0.20) | | | | | | | | | | | | | |
| | **$\beta$-Galactosidase** | APC failed to converge with default parameters | | | | | | | | | | | | 117 |
| | **$\beta$-Glucuronidase** | | | | | | | | | | | | | 85 |
| | **Total** | | | | | | | | | | | | | 202 |
| | (k) APC of the GH3 dataset (SC=0.22) | | | | | | | | | | | | | |
| | **$\beta$-Glucosidase** | 20 | 25 | 61 | 73 | 11 | 10 | 2 | 2 | NA (only 8 clusters) | | | | 204 |
| | **1,4-$\beta$-Xylosidase** | 8 | 7 | 2 | 0 | 4 | 22 | 40 | 16 | | | | | 99 |
| | **Total** | 28 | 32 | 63 | 73 | 15 | 32 | 42 | 18 | | | | | 303 |
| | (l) APC of the GH5 dataset (SC=0.10) | | | | | | | | | | | | | |
| | **Cellulase** | 49 | 55 | 68 | 54 | 37 | 11 | 68 | NA (only 7 clusters) | | | | | 342 |
| | **Endo-$\beta$-Mannanase** | 0 | 7 | 35 | 29 | 14 | 3 | 39 | | | | | | 127 |
| | **Total** | 49 | 62 | 103 | 83 | 51 | 14 | 107 | | | | | | 469 |
| 5 | (m) APC of the SNARE dataset (SC=0.06) | | | | | | | | | | | | | |
| | **non-SNARE** | 28 | 5 | 28 | NA (only 3 clusters) | | | | | | | | | 61 |
| | **SNARE** | 20 | 37 | 0 | | | | | | | | | | 57 |
| | **Total** | 48 | 42 | 28 | | | | | | | | | | 118 |
| | (n) APC of the GPCR1 (SC=0.05) | | | | | | | | | | | | | |
| | **GPCR** | 497 | 206 | 194 | 208 | 332 | 93 | 23 | 57 | 0 | 55 | 0 | 32 | 1697 |
| | **non-GPCR(TM)** | 138 | 54 | 202 | 41 | 71 | 105 | 257 | 445 | 62 | 99 | 176 | 257 | 1907 |
| | **Total** | 635 | 260 | 396 | 249 | 403 | 198 | 280 | 502 | 62 | 154 | 176 | 289 | 3604 |
| | (o) APC of the GPCR2 dataset (SC=0.11) | | | | | | | | | | | | | |
| | **GPCR** | 242 | 668 | 746 | 15 | 26 | NA (only 5 clusters) | | | | | | | 1697 |
| | **non-GPCR(no TM)** | 41 | 77 | 8 | 811 | 924 | | | | | | | | 1861 |
| | **Total** | 283 | 745 | 754 | 826 | 950 | | | | | | | | 3558 |

This table presents the contingency matrices of affinity propagation clustering (APC) for all the datasets present in Groups 4 and 5. A visual representation of the corresponding datapoint distributions in reduced-dimensional space is shown in Figs 15 and 16.

### Runtimes of clustering

We computed the runtime of selected algorithms with respect to (i) the number of clusters, $k$ (Fig 5) and (ii) sample size i.e., total number of protein sequences in the dataset (Fig 6). These were computed on a laptop with an AMD Ryzen 7 4800H processor and 16.0 GB RAM. For each $k$, the runtime was calculated as the sum of runtimes across all datasets. We observe that k-means consistently shows the lowest runtime, followed by agglomerative, Gaussian mixture model (GMM), and spectral clustering. With the exception of GMM, the algorithms exhibit nearly constant runtime across different $k$ values, suggesting that the runtime is independent of $k$.

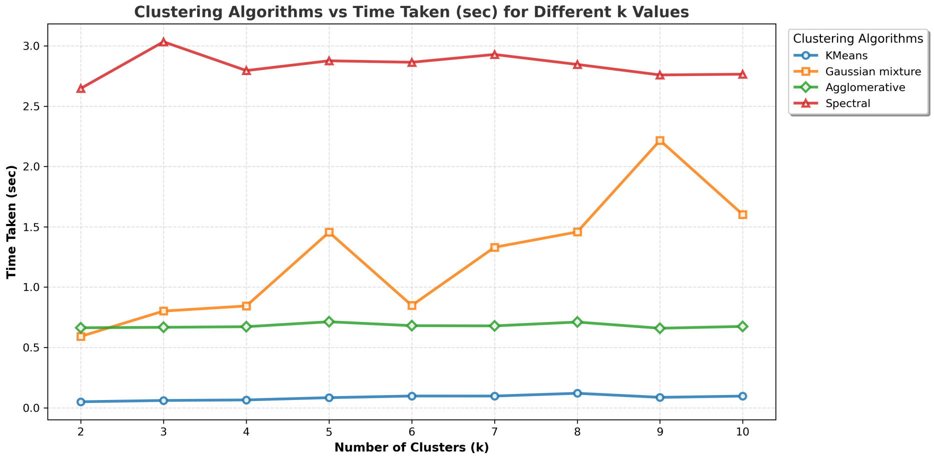


**Fig 5. Runtime comparison of clustering algorithms across different numbers of clusters.** Line plot showing the runtime of four clustering algorithms: k-means, Gaussian mixture model (GMM), agglomerative, and spectral clustering as the number of clusters ($k$) increases from 2 to 10. For each $k$, the runtime was computed as the sum of runtimes across all datasets. Each line represents one clustering algorithm.

The variation runtime as a function of sample size is shown in Fig 6. Here, for each dataset (i.e., sample size), the runtime was computed by summing runtimes over $k = 2$ to 10. In this setting, the runtime of all algorithms increases with sample size. Runtime is the lowest for k-means and the highest for spectral clustering. The largest dataset has 3500 sequences. Overall, it is to be noted that the maximum runtime remained below 13 seconds irrespective of the algorithm, cumulative dataset size, and $k$, even on a laptop.

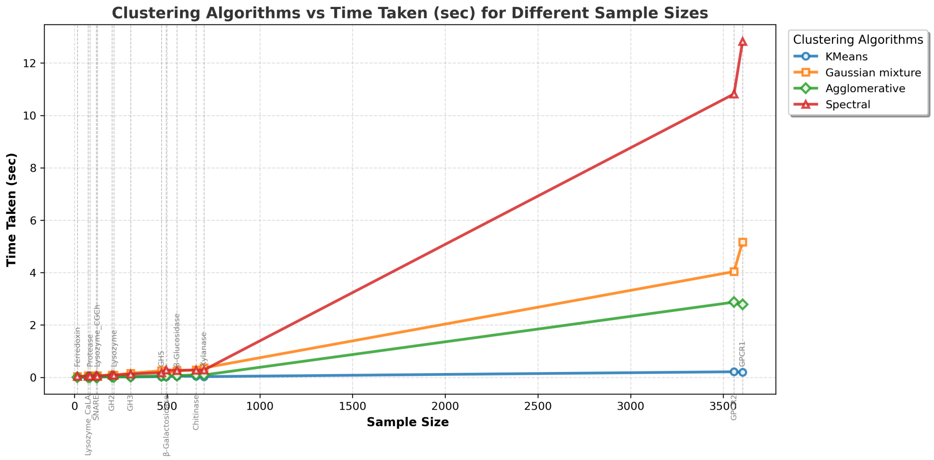


**Fig 6. Runtime comparison of clustering algorithms across datasets of varying sample sizes.** The line plot compares four clustering algorithms: k-means, Gaussian mixture model (GMM), agglomerative, and spectral clustering across different sample sizes. For each sample size(dataset), the total runtime was computed by summing the runtimes for $k = 2$ to 10. Each line represents an algorithm.

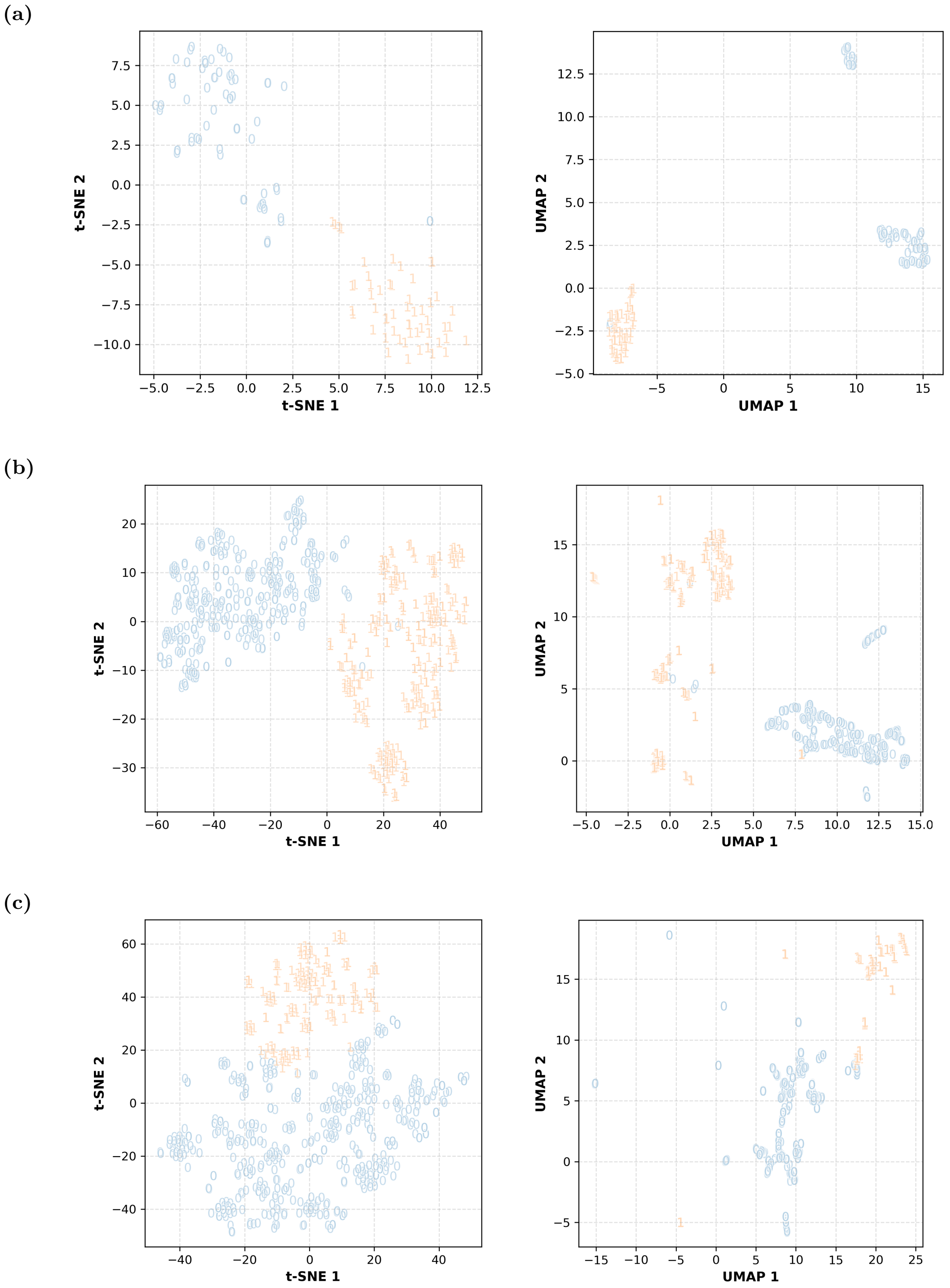


**Fig 7. Distribution of datapoints in reduced-dimensional space for selected clustering results.** The figure shows the distribution of datapoints in various clusters corresponding to the results in Table 4: (a) Lysozyme CGCh; agglomerative, (b) Xyalanse; agglomerative, and (c) Chitinase; spectral. Cluster labels are represented by symbols in the graphs. Both t-SNE and UMAP algorithms have been used for dimensionality reduction.

**(a)**

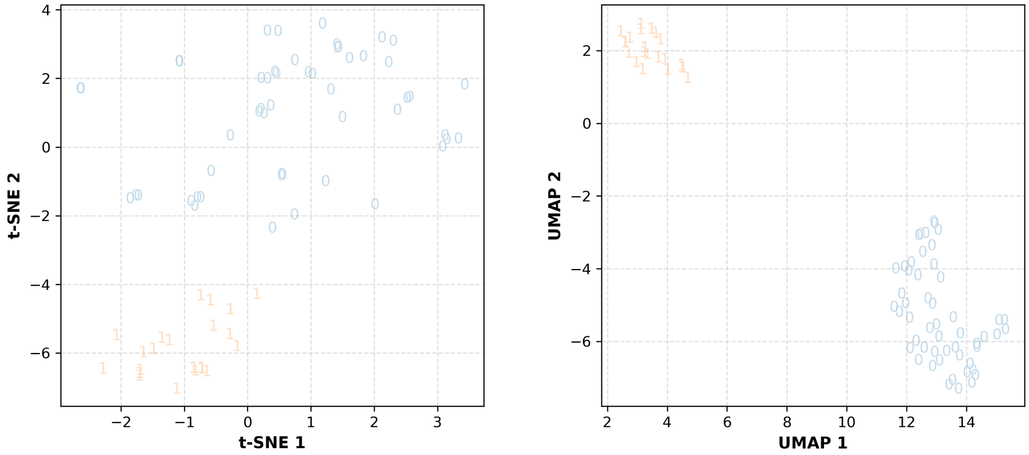


**(b)**

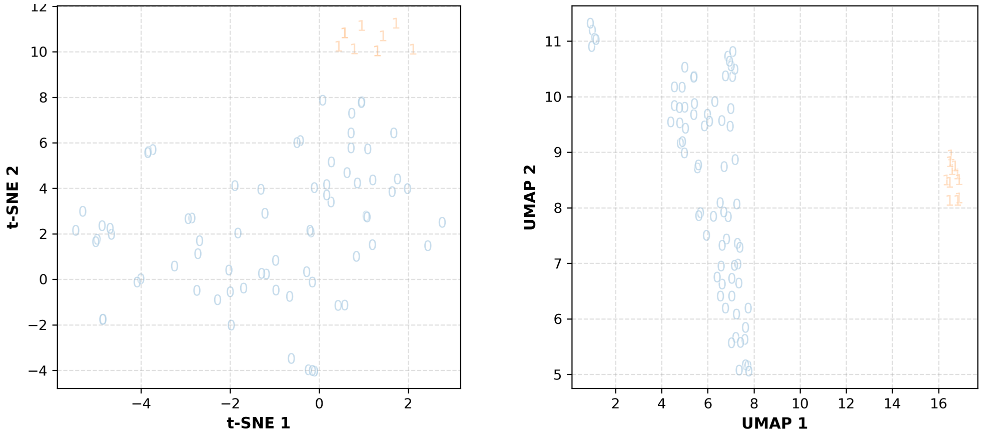


**(c)**

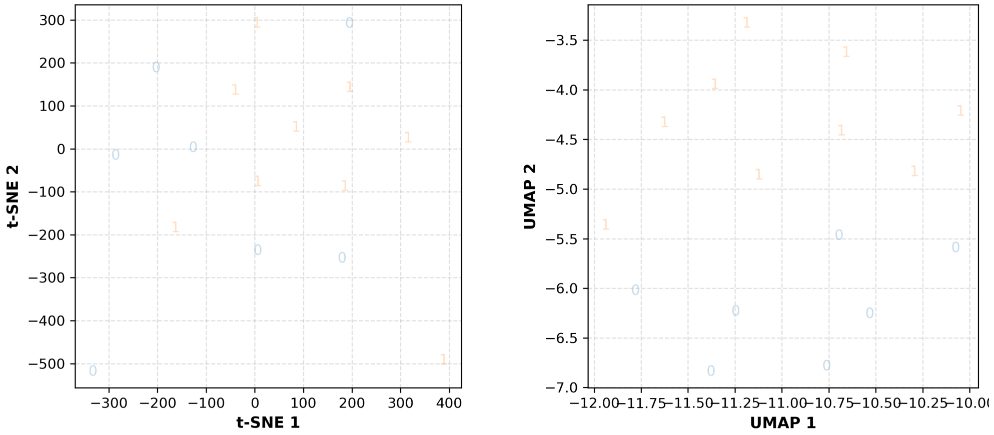


**Fig 8. Distribution of datapoints in reduced-dimensional space for selected clustering results.** The figure shows the distribution of datapoints in various clusters corresponding to the results in Table 4: (a) Lysozyme CaLA; agglomerative, (b) Protease; agglomerative, and (c) Ferredoxin; spectral. Cluster labels are represented by symbols in the graphs. Both t-SNE and UMAP algorithms have been used for dimensionality reduction.

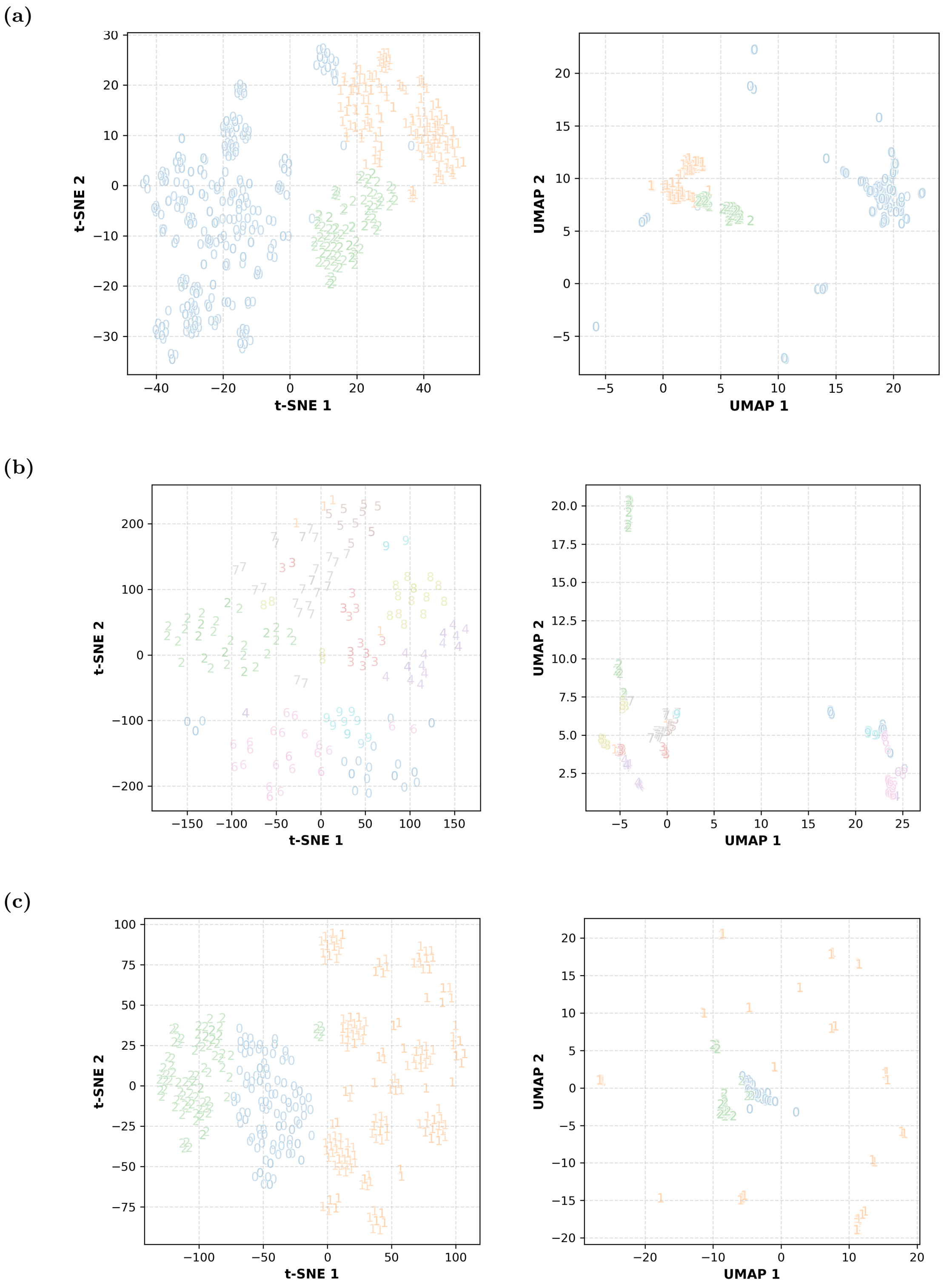


**Fig 9. Distribution of datapoints in reduced-dimensional space for selected clustering results.** The figure shows the distribution of datapoints in various clusters corresponding to the results in Table 4: (a) $\beta$-Glucosidase; agglomerative, (b) Lysozyme; k-means, and (c) $\beta$-Galactosidase; GMM. Cluster labels are represented by symbols in the graphs. Both t-SNE and UMAP algorithms have been used for dimensionality reduction.

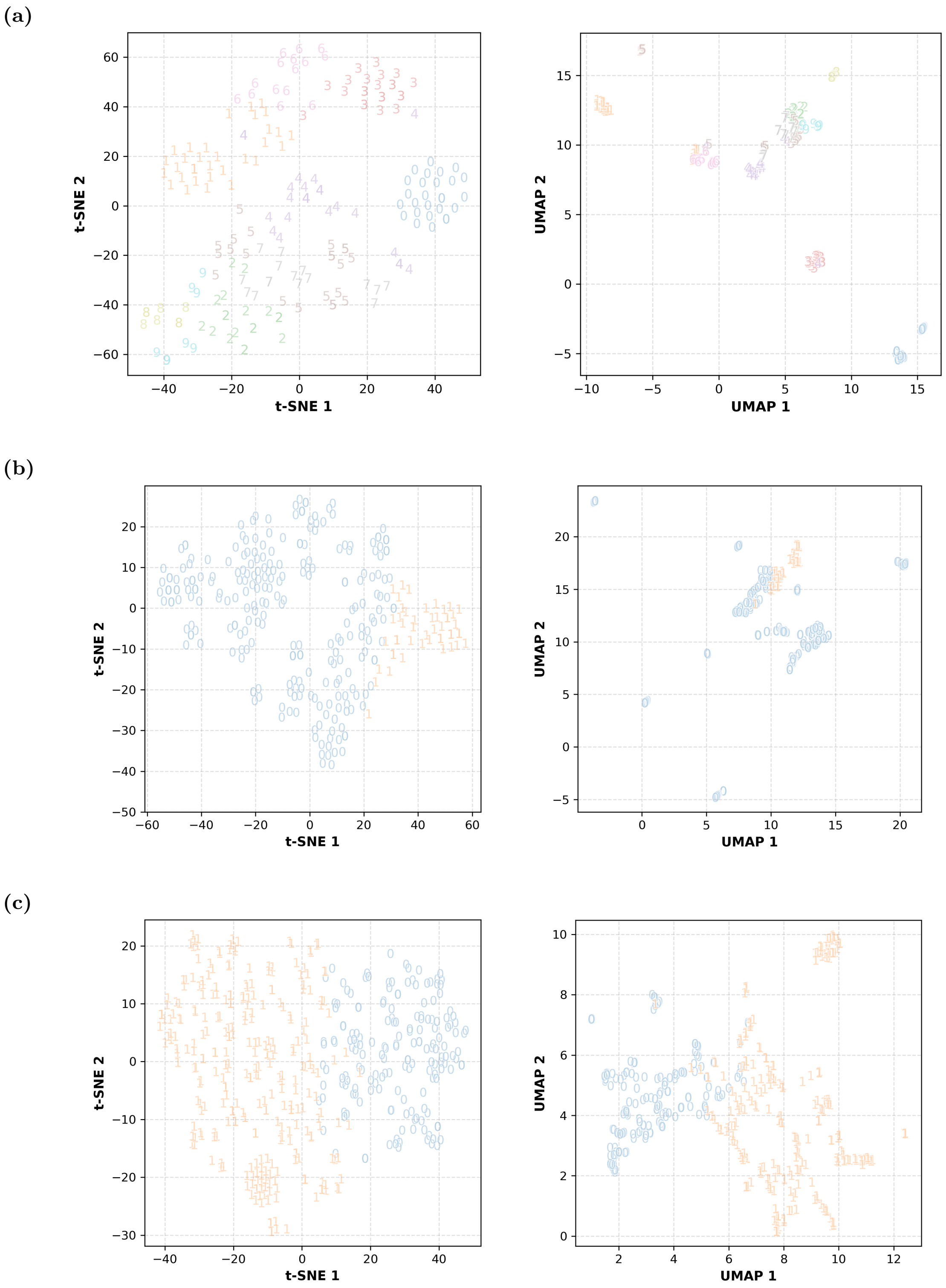


**Fig 10. Distribution of datapoints in reduced-dimensional space for selected clustering results.** The figure shows the distribution of datapoints in various clusters corresponding to the results in Table 5: (a) GH2; k-means, (b) GH3; spectral, and (c) GH5; spectral. Cluster labels are represented by symbols in the graphs. Both t-SNE and UMAP algorithms have been used for dimensionality reduction.

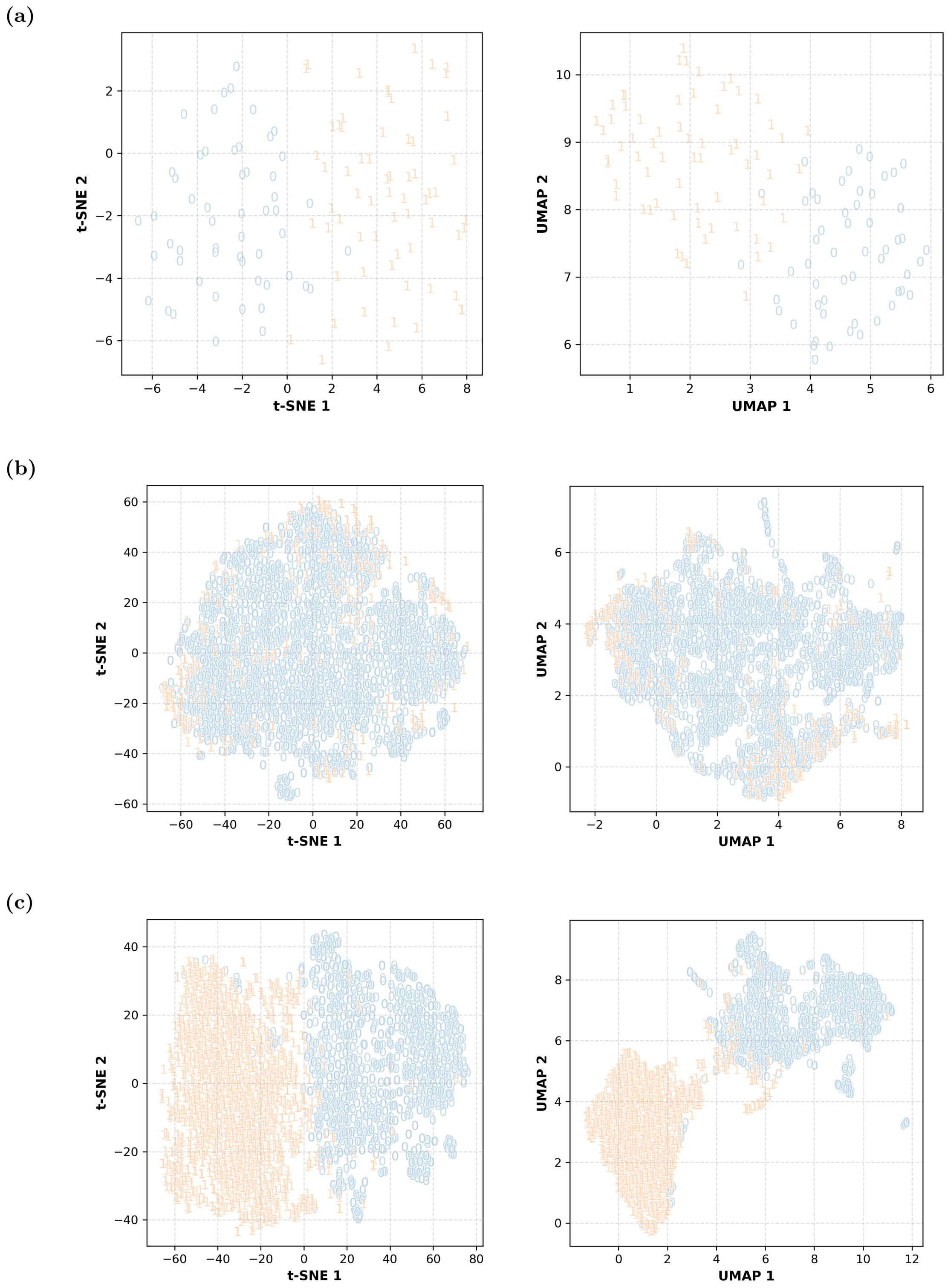


**Fig 11. Distribution of datapoints in reduced-dimensional space for selected clustering results.** The figure shows the distribution of datapoints in various clusters corresponding to the results in Table 5: (a) SNARE; k-means, (b) GPCR1; GMM, and (c) GPCR2; agglomerative. Cluster labels are represented by symbols in the graphs. Both t-SNE and UMAP algorithms have been used for dimensionality reduction.

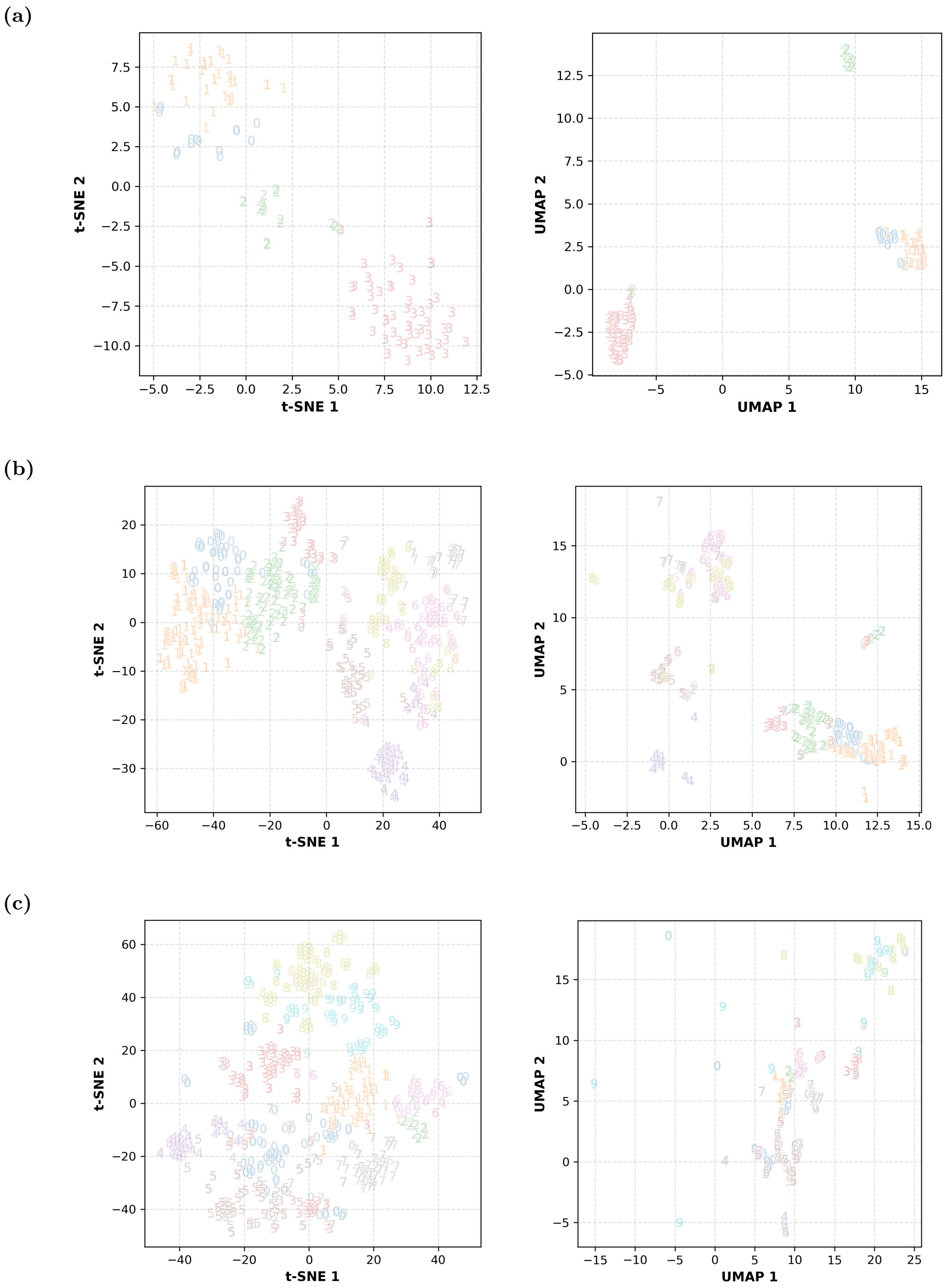


**Fig 12. Distribution of datapoints in reduced-dimensional space for affinity propagation clustering results.** The figure shows the distribution of datapoints in various clusters corresponding to the results in Table 6: APC of (a) Lysozyme CGCh, (b) Xylanase, and (c) Chitinase. Cluster labels are represented by symbols in the graphs. Both t-SNE and UMAP algorithms have been used for dimensionality reduction.

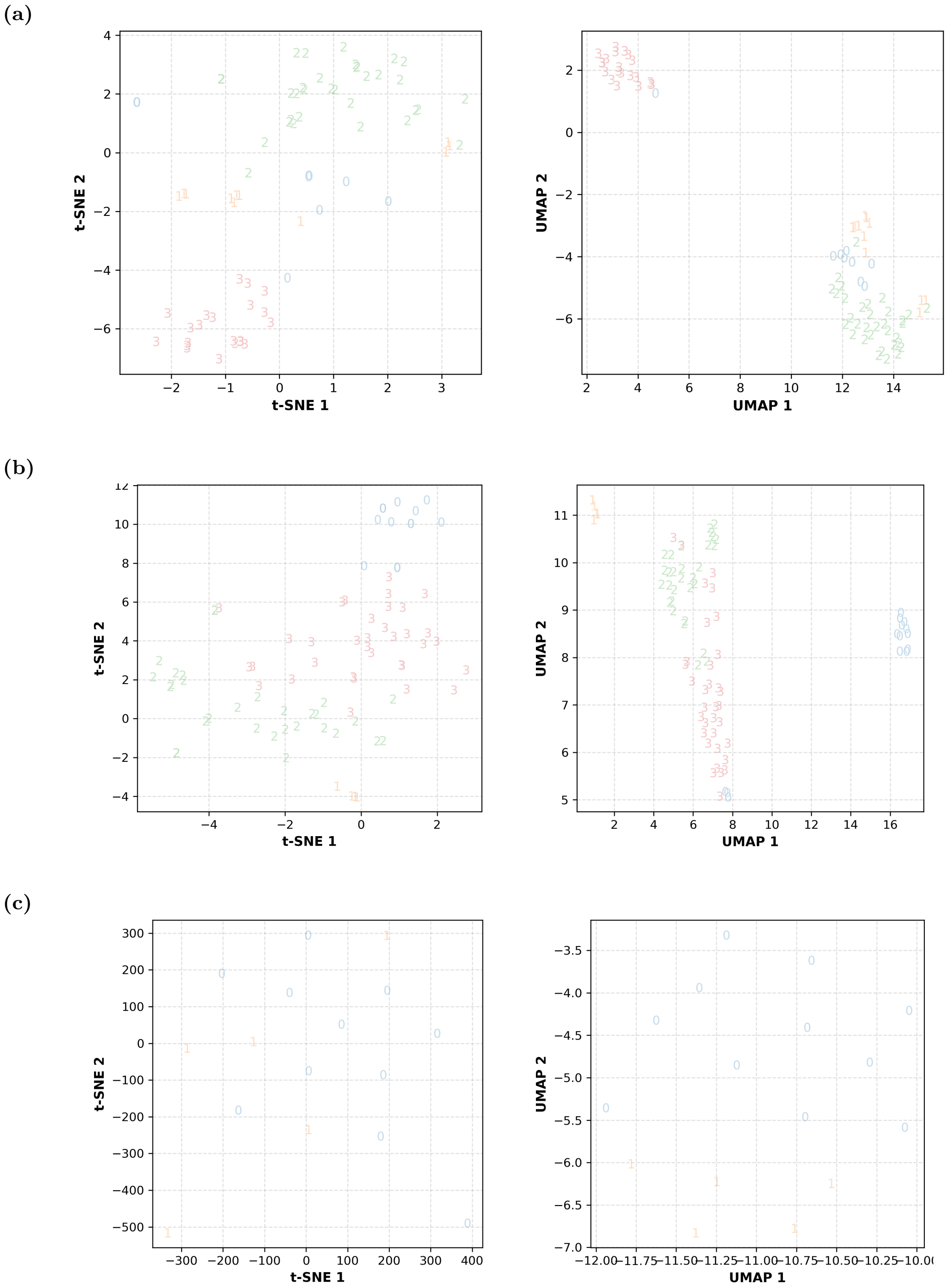


**Fig 13. Distribution of datapoints in reduced-dimensional space for affinity propagation clustering results.** The figure shows the distribution of datapoints in various clusters corresponding to the results in Table 6: APC of (a) Lysozyme CaLA, (b) Protease, and (c) Ferredoxin. Cluster labels are represented by symbols in the graphs. Both t-SNE and UMAP algorithms have been used for dimensionality reduction.

(a)

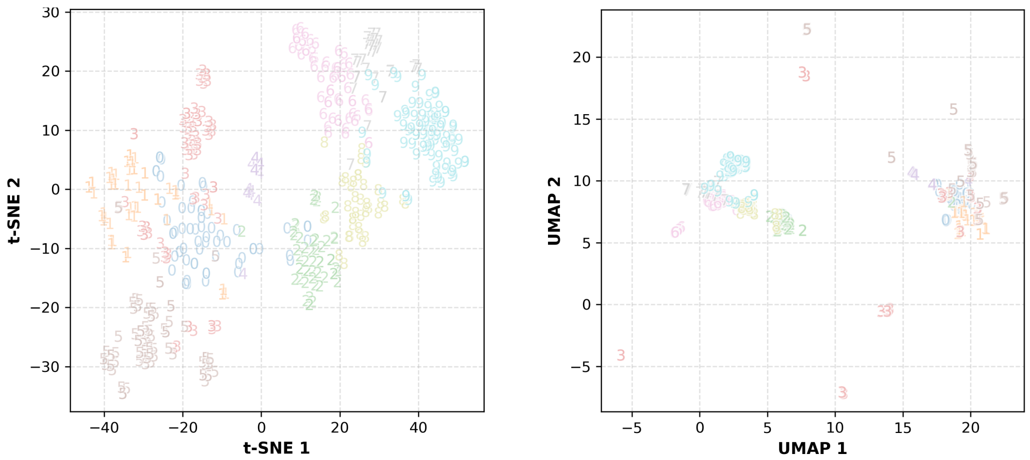


(b)

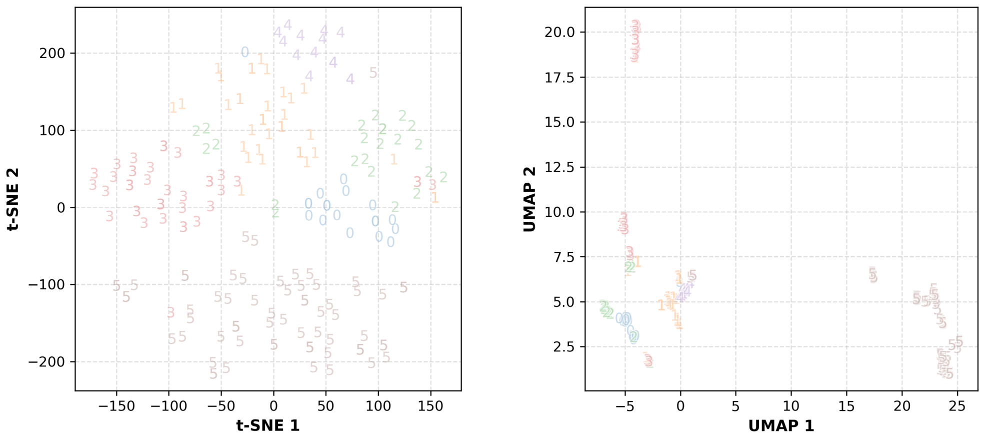


**Fig 14. Distribution of datapoints in reduced-dimensional space for affinity propagation clustering results.** The figure shows the distribution of datapoints in various clusters corresponding to the results in Table 6: APC of (a) $\beta$-Glucosidase, and (b) Lysozyme. $\beta$-Galactosidase is not included due to its inability to converge with default parameters. Cluster labels are represented by symbols in the graphs. Both t-SNE and UMAP algorithms have been used for dimensionality reduction.

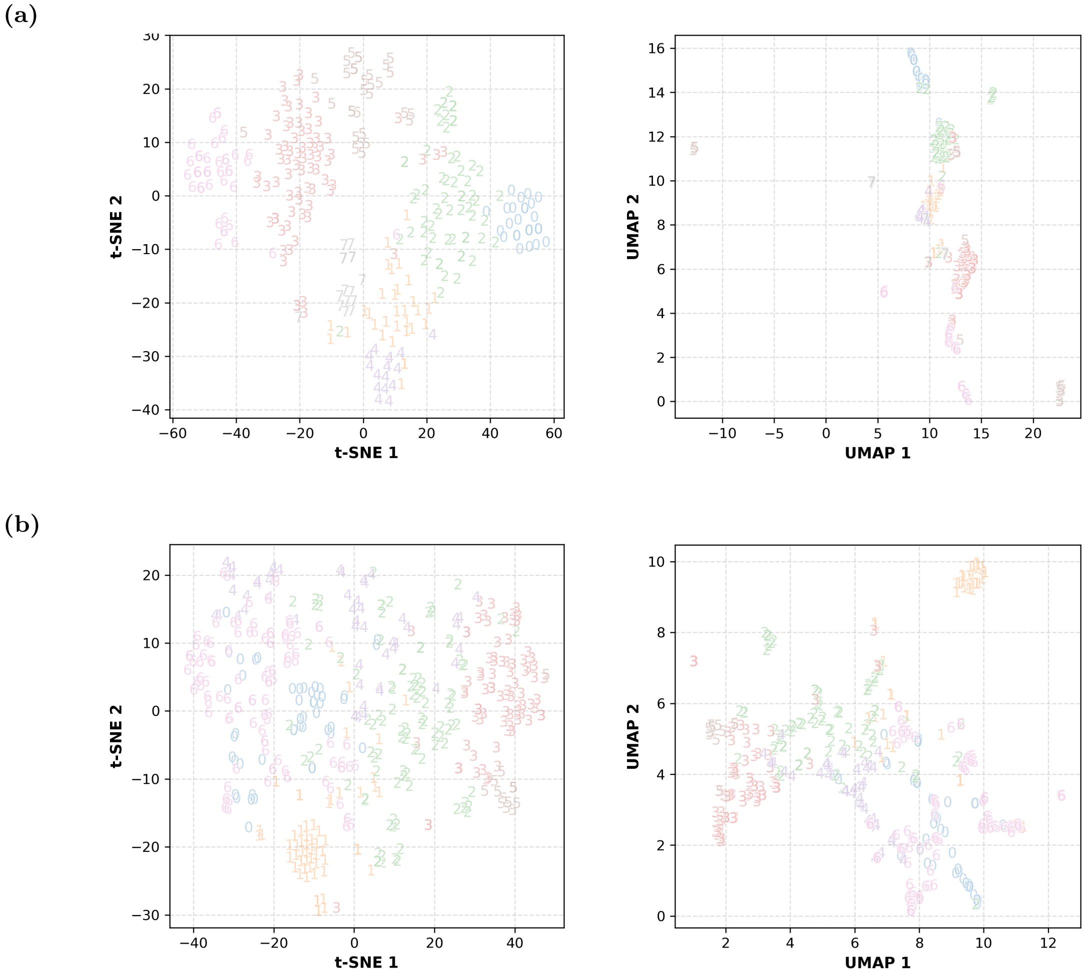


**Fig 15. Distribution of datapoints in reduced-dimensional space for affinity propagation clustering results.** The figure shows the distribution of datapoints in various clusters corresponding to the results in Table 7: APC of (a) GH3, and (b) GH5. GH2 is not included due to its inability to converge with default parameters. Cluster labels are represented by symbols in the graphs. Both t-SNE and UMAP algorithms have been used for dimensionality reduction.

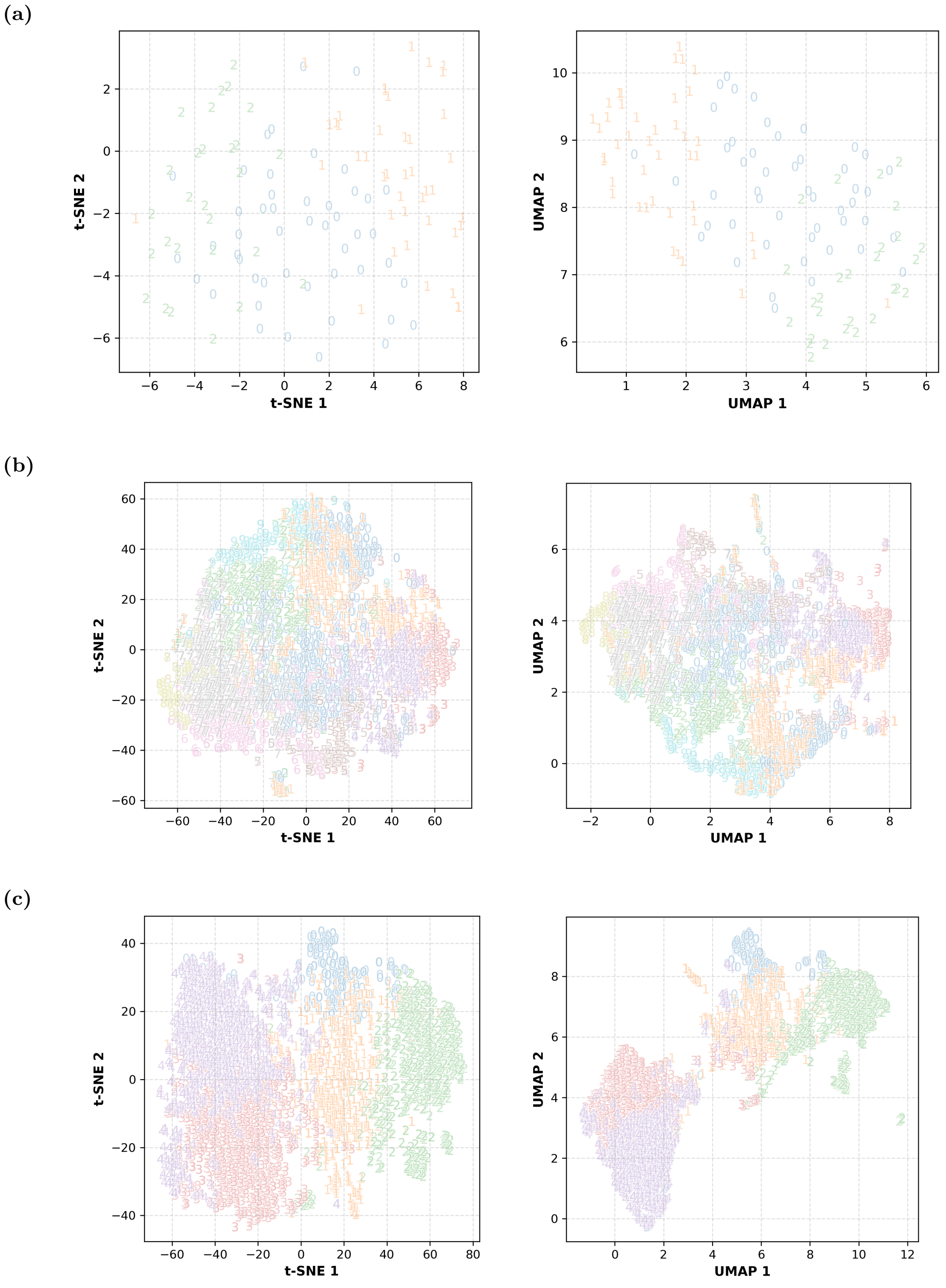


**Fig 16. Distribution of datapoints in reduced-dimensional space for affinity propagation clustering results.** The figure shows the distribution of datapoints in various clusters corresponding to the results in Table 7: APC of (a) SNARE, (b) GPCR1, and (c) GPCR2. Cluster labels are represented by symbols in the graphs. Both t-SNE and UMAP algorithms have been used for dimensionality reduction.
